# Bone regulates locomotion by sustaining motoneuronal mitochondrial function

**DOI:** 10.64898/2026.10.07.750453

**Authors:** David Romeo-Guitart, Tala Tirani, Dimitrije Milunov, Angie K. Torres-Juacida, Boris Lamotte d’Incamps, Carolina Arenas-Plascencia, Sylvère Durand, Ivan Nemazanyy, Stephanie Moriceau, Soham Saha, Gerard Karsenty, Franck Oury

## Abstract

Body-brain communication is crucial for optimal neuronal performance across the lifespan, yet whether peripheral organs regulate spinal cord motoneuron (MN) function through circulating molecules remains unknown. The expression of hormone receptors by spinal MNs raises the possibility that circulating cues allow the spinal cord to sense the body’s energetic state and sustain motor activity. Here, we show that the bone-derived hormones osteocalcin (OCN) signals on spinal MNs via its receptor GPR158. GPR158 is selectively enriched in spinal cholinergic MNs, and its loss, globally or specifically in spinal MNs, or the one of OCN, impairs locomotor function and reduces choline acetyltransferase (ChAT) levels. Mechanistically, OCN signaling through GPR158 fine-tunes autophagy, mitophagy and mitochondrial activity in spinal MNs. Restoring any of these processes, by pharmacologically inducing mitophagy or chemogenetically activating mitochondria, is sufficient to rescue the locomotor deficits of *Gpr158*-deficient mice. Finally, OCN restoration in aged mice reverses age-related locomotor decline through a mechanism that requires MN autophagy and mitophagy. This study reveals that through OCN, the skeleton influences motor functions by fine-tuning motoneuronal mitochondria, and suggests that age-related motor decline is, at least in part, reversible.

## Introduction

Neuronal functions rely on a continuous interplay between peripheral organs and the central nervous system. While body-brain crosstalk has been shown to be essential for maintaining brain fitness, including cognition and metabolic regulation(Che et al., 2026; Oury et al., 2013; Sahasrabudhe et al., 2025; Salvador et al., 2026), its impact on spinal cord function and motoneurons (MNs) remains largely unexplored. Yet the spinal cord sits at the interface between systemic physiology and movement: it integrates descending commands, sensory feedback, and the organism’s metabolic state to control locomotion. Spinal cord motoneurons are critical for translating neural signals into muscle contraction, enabling voluntary and involuntary movements. These neurons are particularly vulnerable to aging due to their high energetic demand and long axonal projections. Human aging is characterized by MN vulnerability, neuromuscular junction dysfunction and denervation (Geertsen et al., 2017; Ham & Rüegg, 2018; Larsson et al., 2019; Sarto et al., 2024), which lead to coordination and strength problems. This raises the possibility that the spinal cord may, like the brain, be a target of hormonal signals that allow the body to sense its energy status and adapt motor output accordingly. Furthermore, understanding whether peripheral organs can influence spinal cord activity through the production and secretion of hormones could reveal the mechanisms by which the locomotor system, composed of the skeleton, nervous system and muscle, maintains its optimal functionality across the lifespan.

Autophagy, the cellular process by which intracellular material is sequestered into autophagosomes and degraded, is essential for integrating hormonal signaling in brain neurons(Blanchet et al., 2025; Glatigny et al., 2019; Moigneu et al., 2023; Rivagorda et al., 2025). In this way, spinal cord autophagy is crucial for MN maintenance, as mice deficient for the key autophagy genes *Atg5* or *Atg7* display MN degeneration and motor deficits. Moreover, underscoring the importance of autophagy in MN function and homeostasis, autophagy induction increases MN regeneration after neurotrauma (Romeo-Guitart et al., 2019, 2020, Hara et al., 2006; Komatsu et al., 2006). However, the upstream regulatory mechanisms of autophagy and its precise role in adult MNs remain unknown.

Mitochondrial homeostasis is particularly critical for spinal cord MNs due to their high energetic requirements. Because MNs have the longest axons in the body and require optimal mitochondrial function to sustain synaptic transmission and excitability along the axon, they are particularly susceptible to the accumulation of dysfunctional mitochondria during aging or stress (Cheng et al., 2022; Misgeld & Schwarz, 2017; Sheng, 2017; Yang et al., 2023). If not effectively cleared through mitophagy, a selective form of autophagy targeting damaged mitochondria, this accumulation can trigger neuronal death, neurodegeneration, and motoneuronal activity deficits during aging (Rygiel et al., 2014).

Traditionally considered solely a structural organ, the bone is now recognized as an endocrine organ, producing hormones that regulate essential physiological functions such as glucose homeostasis, fertility, muscle function during exercise, cognition and the initiation of the acute stress response (Berger et al., 2019; Karsenty & Olson, 2016; Lee et al., 2007; Mera, Laue, Ferron, et al., 2016; Millar et al., 2019; Oury et al., 2013). Osteocalcin (OCN) has risen as a main mediator of bone endocrine effects; OCN mediates energy expenditure and danger response, which are essential neuronal functions (Berger et al., 2019; Khrimian et al., 2017; Lee et al., 2007). To date, the potential influence of hormones, in particular those derived from bone, on spinal cord MNs and motor functions has not yet been investigated.

This study began with a simple exploration to investigate whether hormones may influence locomotor function through spinal cord motoneurons. We observed that MNs express *Gpr158*, and they respond to OCN via this receptor. Mechanistically, OCN–GPR158 coupling promotes mitochondrial function while simultaneously endorsing its selective degradation through mitophagy. This axis is essential, at least in part, for locomotor resilience. These findings identify a new paradigm in body-spinal cord communication and expand the anti-aging benefits of blood-borne rejuvenating factors such as OCN.

## Results

### A GPCR analysis identifies GPR158 as a motoneuron-enriched receptor

We reasoned that, if peripheral organs regulate locomotor functions through the spinal cord, spinal MNs should express receptors able to detect circulating signals. To identify such receptors, we took advantage of published single-nucleus RNA sequencing (snRNAseq) of adult mouse spinal cord (Alkaslasi et al., 2021; Yadav et al., 2023), which resolves MNs into their main subtypes (alpha, gamma, skeletal and visceral) alongside interneurons and glial populations. Since G protein-coupled receptors (GPCRs) constitute the largest family of hormone receptors, we focused our analysis on them.

We built a gene-interaction network for every population and measured how much each GPCR contributes to the integrity of that network, in other words, a way of estimating which receptors act as essential signaling hubs. Focusing on cell-type-specific marker genes, which capture the receptors that distinguish each population rather than those shared by all cells, we found that alpha MNs are the spinal cell type whose GPCRs have the greatest impact on network integrity (**Fig. 1a and S1b**). This indicates that alpha MNs depend on GPCR signaling to an unusually high degree. To ask which types of receptors drive this dependence, we ranked all GPCRs by their impact on network integrity and divided them into five equal groups, or bins, from the highest-impact receptors (Bin 1) to the lowest (Bin 5). We then examined the family composition of each bin, which showed that high- and low-impact receptors belonged to distinct GPCR families (**Fig. 1b**).

**Figure 1:**
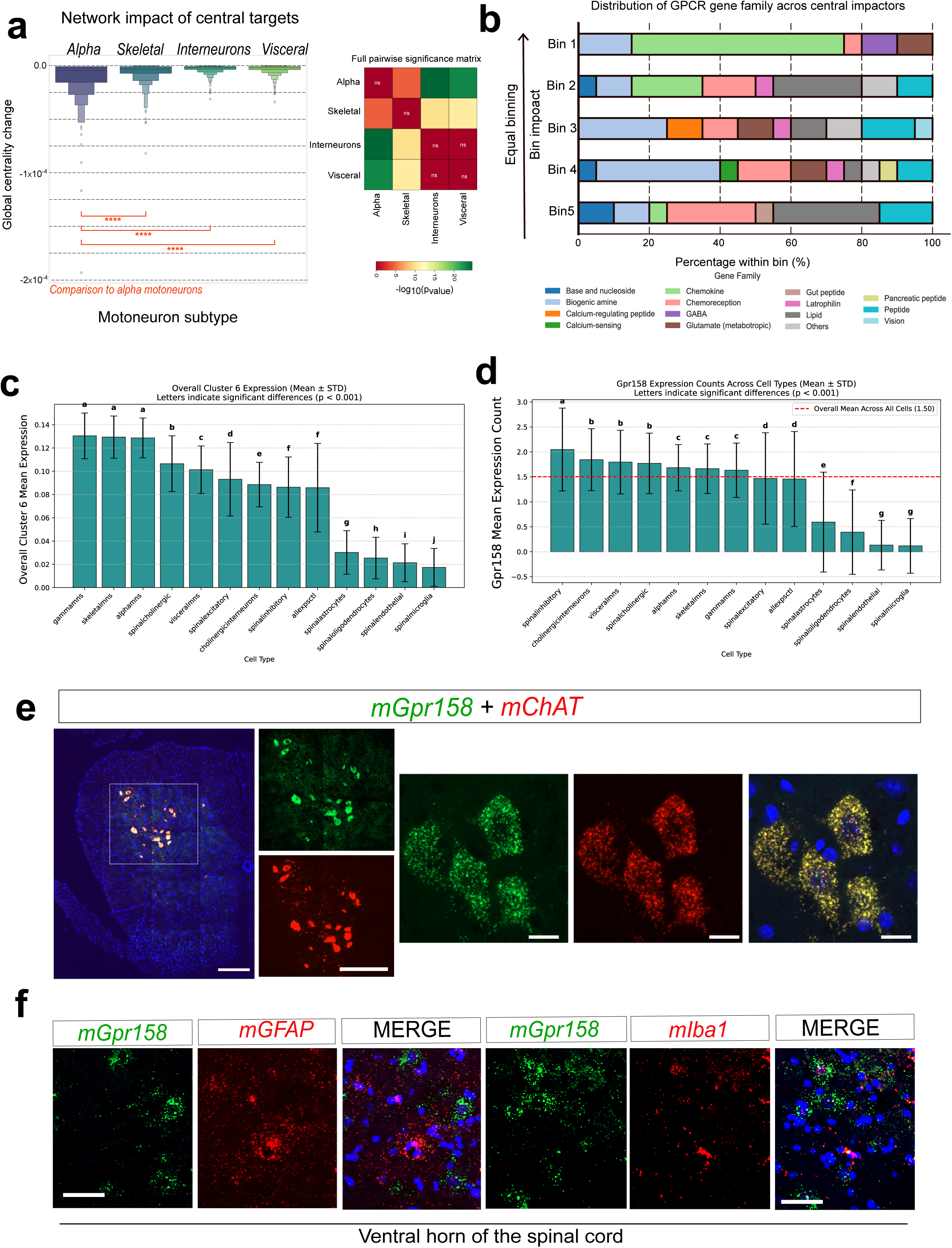
GPR158 is a motoneuron-enriched hormonal receptor in the spinal cord. **a.-** Network impact analysis of GPCRs across spinal cell types, based on cell-type-specific marker genes. Alpha motoneurons (MNs) show the greatest GPCR impact on network integrity, indicating a strong dependence on GPCR signaling. **b.-** GPCRs ranked by network impact and divided into five equal bins (Bin 1, highest; Bin 5, lowest), showing family composition per bin. **c.-** Mean expression of the MN-enriched GPCR co-expression module (Cluster 6) across spinal cord cell types. Alpha, skeletal and gamma MNs show the highest expression; letters indicate statistically indistinguishable groups. **d.-** *Gpr158* expression across spinal cell types. Gpr158 is enriched in neuronal populations, including MNs, and low or absent in astrocytes, oligodendrocytes, endothelial cells and microglia. Dashed line indicates the overall mean across all cells. **e.-** RNAscope *in situ* hybridisation in the spinal cord using *mGpr158* probes (green) with *mChat* probes to stain cholinergic neurons (mainly MNs) (red); nuclei counterstained with DAPI (blue). Boxed region is magnified on the right. Scale bar = 200 and 20 µm. **f.-** RNAscope *in situ* hybridisation in the spinal cord ventral horn using *mGpr158* probes (green) with *mIba1* (microglia) or *mGfap* (astroglia) probes counterstained with DAPI (blue). Boxed region is magnified on the right. Scale bar = 50 µm. In panels (**c**) and (**d**), groups sharing the same letter are not significantly different (compact letter display; P < 0.001). P values in (a) were determined by the Mann-Whitney test versus alpha MNs.

We repeated this analysis using all expressed genes to capture the full set of GPCRs present in each population. Hierarchical clustering of GPCR expression across all cell types grouped receptors into co-expression modules, sets of GPCRs that are expressed together in the same populations. One of these modules, Cluster 6, was selectively enriched in MNs (**Fig. S1a**). Quantifying the average expression of this module confirmed that alpha, skeletal and gamma MNs express it at significantly higher levels than any other spinal cell type (**Fig. 1c**). STRING network enrichment analysis of this module was dominated by class C / family 3 GPCRs, the structural class to which GPR158 belongs (**Fig. S1c**). Within this MN-enriched module, we focused on Gpr158, one of the most motor neuron-enriched receptors and a class C GPCR. Examining its expression across cell types confirmed that Gpr158 is restricted to neuronal populations, including MNs, and is low or absent in astrocytes, microglia and oligodendrocytes (**Fig. 1d**). In agreement with these data, qPCR across the motor system detected *Gpr158* in the motor cortex, brainstem and spinal cord, while it was absent from skeletal muscle (**Fig. S1d**). We observed that GPR158 protein is also present in the spinal cord, sciatic nerve and dorsal root ganglia, while absent in gastrocnemius muscle (**Fig. S1e**). To validate this finding at the anatomical level, we performed RNAscope fluorescent *in situ* hybridization on spinal cord sections. *Gpr158* mRNA co-localized extensively with choline acetyltransferase (*ChAT*)-positive MNs in the ventral horn, with no detectable signal in non-neuronal populations such as microglia and astroglia (**Fig. 1e, f**). Of note, Gpr158 signal was completely absent in spinal cord sections from *Gpr158* -/- mice, confirming probe specificity (**Fig. S1f**). Altogether, these results identify Gpr158 as an MN-enriched hormonal receptor that may influence motor function.

### GPR158 in spinal motor neurons is required for locomotor function

To test whether GPR158 expression in spinal cord MNs is required for motor behaviour, we performed 3 complementary experiments. First, we assessed *Gpr158* -/- male and female 3-month-old mice using the accelerating rotarod and beam bar test to assess coordination. Both cohorts showed reduced latency to fall and coordination deficits compared to wild-type littermates (**Fig. 2a, b, and S2d**). To determine whether the motor deficit is intrinsic to the spinal cord, we generated two mouse models: a broader Gpr158 knockdown using intrathecal delivery of an AAV carrying shRNA against *Gpr158* under the H1 promoter (AAVrh10-H1-sh*Gpr158*), and a selective downregulation with the MN-specific miR218 (AAVrh10-miR218-sh*Gpr158)* (Thiebes et al., 2015). Knockdown efficiency was confirmed at both mRNA and protein levels for both models (**Fig. 2c, f and S2a, c**), and selective MN infection was validated for the miR218-promoter with Immunofluorescence against ChAT (**Fig. S2b**). Both approaches recapitulated the locomotor deficits observed by rotarod and beam bar of the global knockout, which indicate that GPR158 regulates locomotor functions through its presence in cholinergic spinal MNs (**Fig. 2d, e, f, h**). Consistent with the fact that the deficit is specific to motor performance and coordination rather than general locomotor ambulatory activity, open field locomotion was unaffected in both models of Gpr158 downregulation (**Fig. S2e**). Together, these results demonstrate that GPR158 is required for normal locomotor function and that this requirement is cell-autonomous to spinal MNs.

**Figure 2:**
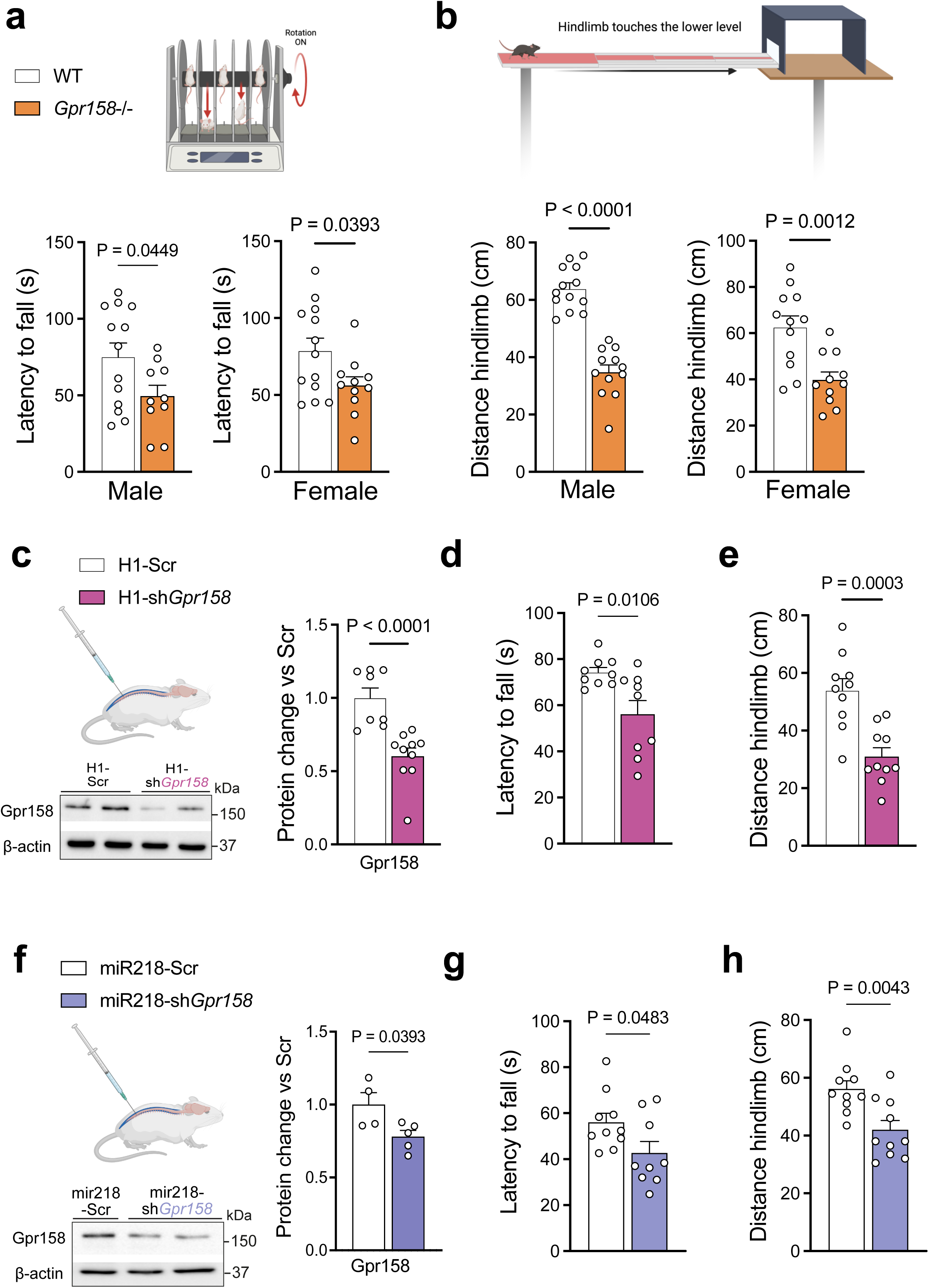
GPR158 in spinal cord motoneurons regulates locomotor functions. **a. b-** Rotarod performance (**a**) and Beam bar coordination (**b**) test in 3-month wild-type (WT) and *Gpr158* -/- male and female mice. **c-** Western blot and quantification 3 weeks after intrathecal injections with AAV-H1-Scramble (Scr) or AAV-H1-shRNA*-shGpr158*, and respective quantification of GPR158 in the spinal cord. **d, e-** Rotarod performance (**d**) and Beam bar coordination test (**e**) performed in mice 3 weeks after intrathecal injections of AAV-H1-shRNA*-*Scramble or *shGpr158.* **f.-** Western blot and quantification 3 weeks after intrathecal injections with AAV-mir218-Scramble (Scr) or AAV-mir218-shRNA*-shGpr158*. **g, h-** Rotarod performance (**g**) and Beam bar coordination test (**h**) performed in mice 3 weeks after intrathecal injections with AAV-mir218-shRNA*-*Scramble or *shGpr158.* Data are expressed as mean ± SEM. The P values were determined by two-tailed Student’s t-test (**a**, **b**, **d**, **e**, **f**, **g, h**) or Mann-Whitney test (**c**). Each dot represents an independent mouse (Spinal cord for molecular biology, or behavioral test).

### Osteocalcin signals to spinal motoneurons via GPR158 and maintains motor function

Given prior evidence that osteocalcin signals in the nervous system through GPR158 (Khrimian et al., 2017), we tested whether spinal cord motoneurons respond to the bone-derived hormone OCN by complementary means.

Since OCN regulates GABAergic and glutamatergic tone in the brain, we asked whether it modifies electrical activity in spinal neurons. We performed whole-cell current-clamp recordings in spinal cord slices, identifying MNs by location in the ventral horn and size. Bath application of OCN (10 and 100ng/mL) changed the resting membrane potential of ventral horn neurons: 4 were hyperpolarized (-4.4 to -1.1mV) and 4 were depolarized (+0.4 to +10.6mV). Neuronal excitability, assessed with slow triangular current ramps, changed in accordance with these shifts in membrane potential, with hyperpolarized cells showing reduced maximal firing and depolarized cells showing increased firing (**Fig. S3a, b**). Notably, the hyperpolarized, less excitable cells corresponded to those with the lowest input resistance, i.e., the largest neurons, which include alpha motoneurons and which were inhibited by OCN (**Fig. S3b**). Thus, OCN modulates the electrical properties of spinal neurons with a variable effect across cells.

We next asked whether OCN shapes the cholinergic identity of spinal MNs, by analyzing the essential ChAT and the vesicular acetylcholine transporter (VAChT) (Oury et al., 2013). We found that *Ocn -/-, Gpr158*-/-, and mice in which Gpr158 was knocked down in the spinal cord MNs presented *ChAT* and *VAChT* expression at a lower level in the spinal cord that control mice (**Fig. S3c**). ChAT protein levels was also decreased in the spinal cord of these mice (**Fig. 3a**). To assess whether OCN activates MNs by GPR158, we systemically injected OCN and assessed cFOS positive (cFOS^+^) MNs. OCN induced robust cFOS nuclear accumulation in spinal MNs, and this induction was abolished in *Gpr158*-/-, indicating that OCN activates spinal MNs through GPR158 (**Fig. 3b**). Finally, intrathecal OCN increased ChAT mRNA and protein levels in the spinal cord (**Fig. 3c and S3d**). OCN ALSO requires GPR158 to increase ChAT levels in the spinal cord (**Fig 3d**). Those effects were further validated *in vitro*, by using a MN-cell line like NSC34. OCN treatment increased ChAT protein levels, and this increase was blocked upon sh*Gpr158* transfection (**Fig. 3e and S3e**). Finally, *Ocn*-/- and *Gpr158*-/- mice presented reduced mRNA levels of *BDNF* and *CREB* in the lumbar spinal cord (**Fig. S3f**), a signaling axis previously shown to mediate OCN effects in the brain (Khrimian et al., 2017; Rivagorda et al., 2025). Consistently, *Ocn* -/- male and female mice displayed significant deficits in rotarod performance and beam bar coordination (**Fig. 3f, g, S3g**), closely paralleling the phenotype of GPR158-targeted animals. Altogether, these data indicate that OCN signaling through GPR158 in spinal cord MNs influences locomotor function.

**Figure 3:**
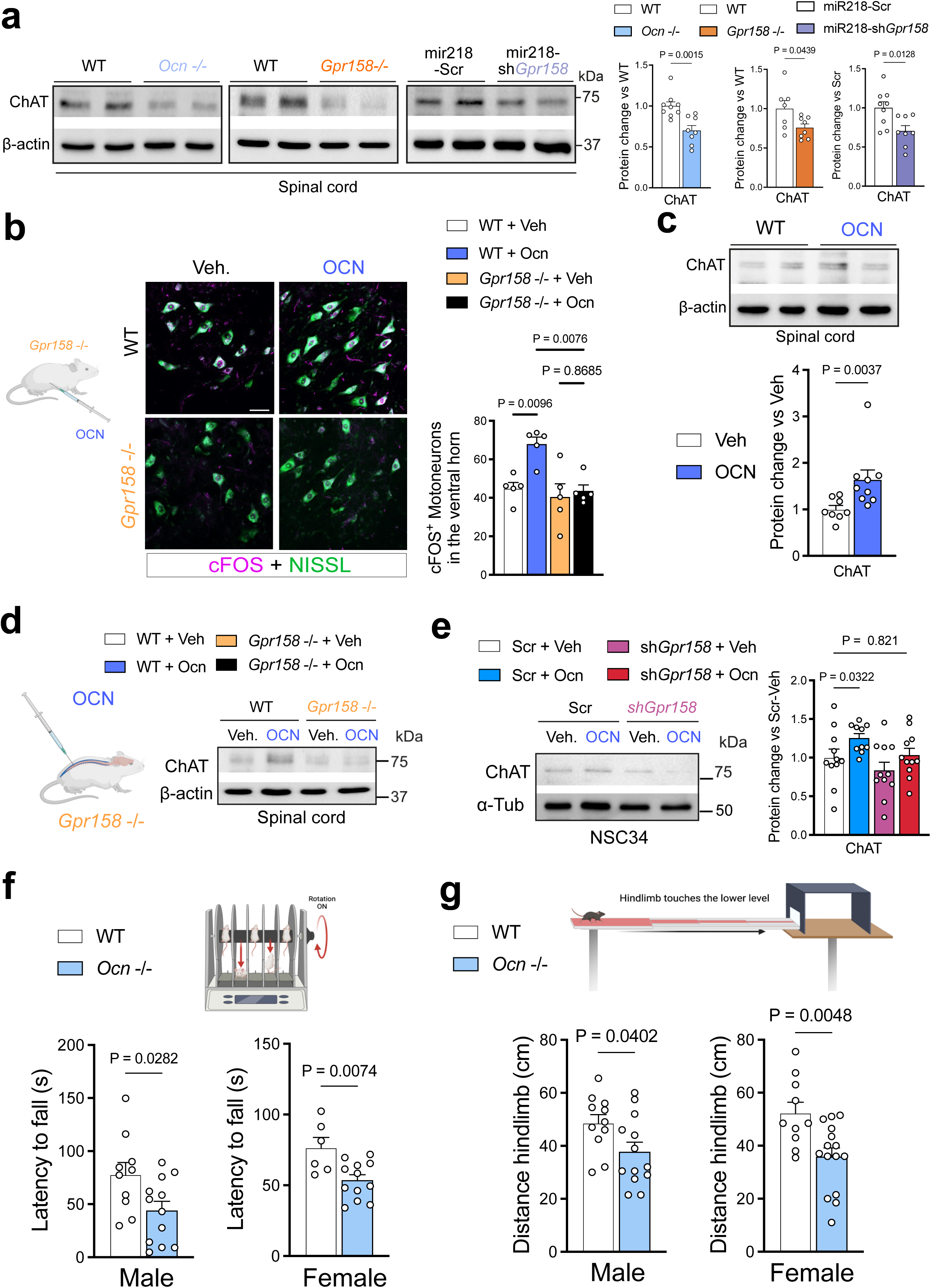
Osteocalcin–GPR158 signaling maintains motoneuronal ChAT and locomotor function. **a.-** Representative western blot and quantification of ChAT in the spinal cord from wild-type WT, *Ocn-/-, Gpr158-/-* and upon the intrathecal injections with AAV-mir218-Scramble (Scr) or AAV-mir218-shRNA*-shGpr158.* **b.-** Immunofluorescence and quantification of cFOS^+^ (Magenta) motoneurons in the ventral horn of the spinal cord of WT and *Gpr158*-/- mice after intraperitoneal injection of OCN. Scale bar = 50 µm**. c.-** Representative western blot and quantification of ChAT in the spinal cord from wild-type mice 4 hours after the intrathecal injection of OCN. **d.-** Representative western blot of ChAT in WT and *Gpr158*-/- mouse spinal cord 4 h after intrathecal injections with vehicle or OCN. **e.-** Representative western blot and quantification analysis of ChAT in the NSC34 cell line transfected with Scramble or sh*Gpr158*, and treated for 1 hour with vehicle or OCN. **f, g-** Rotarod performance (**f**) and Beam bar coordination (**g**) tests in 3-month wild-type (WT) and *Ocn* -/- male and female mice. Data are expressed as mean ± SEM. The P values were determined by Student’s t-test (**a**, **e**, **f**, **g**), Mann-Whitney test (**c**), one-way ANOVA (Holm-Šídák’s multiple comparisons test Dunnett’s multiple comparison test) (**b**). Each dot represents an independent mouse (Spinal cord for molecular biology, or behavioral test) or NSC34 independent preparation.

### Unbiased proteomics identifies autophagy and mitochondrial pathways downstream of OCN–GPR158

To identify the molecular events downstream of OCN signaling in MNs, we performed proteomics on spinal cord from two complementary models: wild-type (WT) mice receiving intrathecal OCN, and Gpr158-/- versus WT mice. Unsupervised hierarchical clustering separated the three groups, indicating that OCN stimulation and GPR158 loss reshape the spinal cord proteome, producing opposite proteomic signatures and indicating that they act on the same axis (**Fig. 4a**). Grouping the proteins by how they behaved across the three conditions separated them into eight clusters (C1–C8), which we further organized into three main categories according to their patterns (**Fig. 4b, S4a**). The average levels of these clusters differed significantly between WT, WT + OCN and *Gpr158* -/- in most cases (clusters C1–C5, C7, and C8). Of the three categories, Category 2 contained the largest share of regulated proteins (47.87%), followed by Category 3 (22.98%) and Category 1 (16.84%), with a smaller group of uncharacterized proteins (12.32%) (**Fig. S4b**).

**Figure 4.**
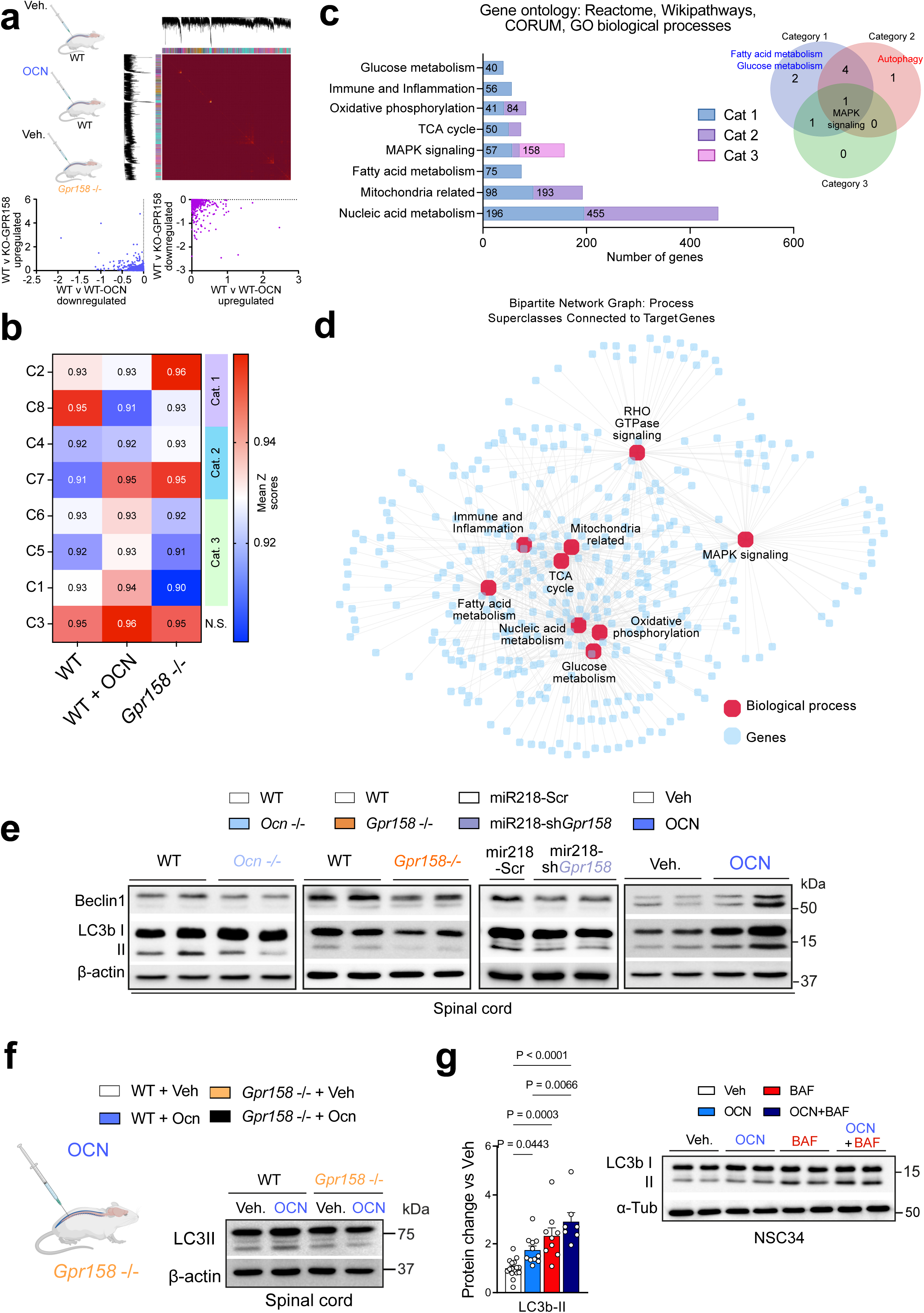
Unbiased proteomics identify OCN/GPR158 axis as a regulator of mitophagy and mitochondrial functions. **a.-** Hierarchical clustering heatmap and scatter plots of the lumbar spinal cord proteome from wild-type (WT) intrathecally injected with Vehicle or OCN, and *Gpr158*-/- intrathecally injected with Veh. **b.-** Mean Z-scores of each cluster across the three conditions; showing that OCN treatment and *Gpr15* -/- spinal cords present protein expression trends in opposite directions. Clusters (C1-C8) are grouped into three statistically significant categories (Cat. 1–3; ANOVA with post hoc correction: Tukey’s test). **c-** Functional enrichment analysis (Reactome, WikiPathways, CORUM, GO biological processes) of the regulated proteins, with the number of proteins per pathway and category. The Venn diagram shows pathways shared between categories**. d.-** Bipartite network linking regulated proteins (blue nodes) to their biological process superclasses (red nodes), showing an interconnected mitochondrial and metabolic core**. e.-** Representative western blot of Beclin1 and LC3b the lumbar spinal cord from wild-type WT, *Ocn-/-, Gpr158-/-,* upon the intrathecal injections with AAV-mir218-Scramble (Scr) or AAV-mir218-shRNA*-shGpr158,* and upon intrathecal injection of Vehicle or OCN. **f.-** Representative western blot of LC3b in WT and *Gpr158*-/- mouse spinal cord 4 h after intrathecal injections with vehicle or OCN. **g.-** Representative western blot and quantification of LC3b-II accumulation in NSC34 cells treated for 4 h with vehicle, OCN, Bafilomycin, or OCN + Bafilomycin. α-Tubulin was used as a loading control for each sample. Data are expressed as mean ± SEM. The P values were determined one-way ANOVA (Tukey’s multiple comparison test) (**g**). Each dot represents an NSC34 independent preparation.

Functional enrichment analysis (Reactome, WikiPathways, CORUM, and GO biological processes) showed that the regulated proteins belonged mainly to metabolic and signaling pathways. The most enriched were nucleic acid metabolism (455 proteins in Category 3; 196 in Categories 1 and 2), mitochondria-related processes (193; 98), MAPK signaling (158; 57), oxidative phosphorylation (84; 41), fatty acid metabolism (75), immune and inflammation pathways (56), tricarboxylic acid (TCA) cycle (50) and glucose metabolism (40). Comparing the categories, they showed both shared and category-specific pathways, with autophagy, mitophagy, MAPK signaling and fatty acid metabolism standing out (**Fig. 4c**).

To assess how these pathways connect to each other, we built a network linking each regulated protein to the biological processes it belongs to (**Fig. 4d**). In this network, the large red dots are the main biological processes and the small blue dots are the individual proteins, with lines showing which protein belongs to which process. The network revealed that the metabolic and mitochondrial processes, glucose metabolism, oxidative phosphorylation, TCA cycle, and mitochondria-related functions, were densely interconnected, forming a tightly linked metabolic core, while nucleic acid metabolism and MAPK signaling engaged broader sets of proteins loss (**Fig. 4d**).

Two additional observations support OCN-mediated mitochondrial recycling. First, a subset of mitochondrial proteins was selectively reduced in OCN-treated spinal cords (**Fig. S4c**). This is consistent with mitophagy induction, as degradation of mitochondria would be expected to reduce the abundance of mitochondrial proteins. Second, ranking all proteins by both their degree of regulation and their centrality within the network identified the core autophagy machinery (Beclin1, Atg13 and Gabarapl1) as the most central and most strongly regulated nodes (**Fig. S4d**). Together, these analyses positioned autophagy, and Beclin1 in particular, as a central effector of the OCN–GPR158 axis, which we next tested directly.

The data presented above prompted us to test to whether OCN directly controls autophagy machinery in the spinal cord. Ocn-/- and Gpr158-/- mice, as well as MN-specific Gpr158 knockdown, showed reduced LC3b-II and Beclin1 protein levels (autophagosome content and autophagy initiation markers, respectively), indicating less autophagy machinery (**Fig. 4e and S4f**). Conversely, intrathecal OCN injection increased Beclin-1 and LC3b-II levels in the spinal cord, as detected by western blot, indicating an accumulation of autophagosomes. Importantly, intrathecal OCN failed to increase LC3b-II in Gpr158-/- spinal cord (**Fig, 4f**). To resolve this at cellular resolution and distinguish autophagy induction from impaired degradation, we used NSC34 cells. OCN treatment increased LC3b-II, and this increase was further enhanced by bafilomycin A1 (BAF), which blocks lysosomal degradation, indicating that OCN promotes autophagosome formation (**Fig. 4g**). Together, these results establish that the OCN signaling through GPR158 in spinal MNs controls the autophagy machinery, and suggest that OCN may influence locomotor functions by fine-tuning autophagy and controlling mitochondrial quality control.

### Autophagy and mitophagy manipulation rescue motor deficits in GPR158-deficient mice

We reasoned that if the locomotor deficit in Gpr158-/- mice is caused at least in part by reduced autophagy, restoring autophagic machinery should rescue the phenotype. Thus, we first administered TETA, a caloric restriction mimetic that activates autophagy to Gpr158-/- mice (Castoldi et al., 2020; Pietrocola et al., 2020). Systemic TETA significantly improved rotarod and beam bar performance in Gpr158 -/- mice, indicating that systemic autophagy restores locomotor functions (**Fig. 5a and S5a**). Consistently, intrathecal TAT-Beclin1, a cell-permeable autophagy-inducing peptide, also rescued rotarod in Gpr158 -/- mice (**Fig. 5b**), confirming that pharmacological activation of autophagy is sufficient to bypass the loss of OCN signaling. To test whether autophagy is required for motor function, we inhibited it through complementary approaches. We intrathecally injected an AAVrh10-SYN1-shBeclin1 to selectively downregulate autophagy in spinal cord neurons. First, we observed the efficient downregulation of Beclin1 at mRNA and protein levels, which was accompanied by less LC3b-II and ChAT (**Fig. 5c, S5b, c**). Those changes were accompanied by impaired rotarod performance without affecting Open Field locomotion (**Fig. 5c and S5c**). Pharmacological inhibition of autophagy with intrathecal leupeptin or spautin-1 similarly induced motor deficits in wild-type mice (**Fig**. **5d**). Hence, basal autophagy in spinal MNs is necessary to maintain normal motor function.

**Figure 5:**
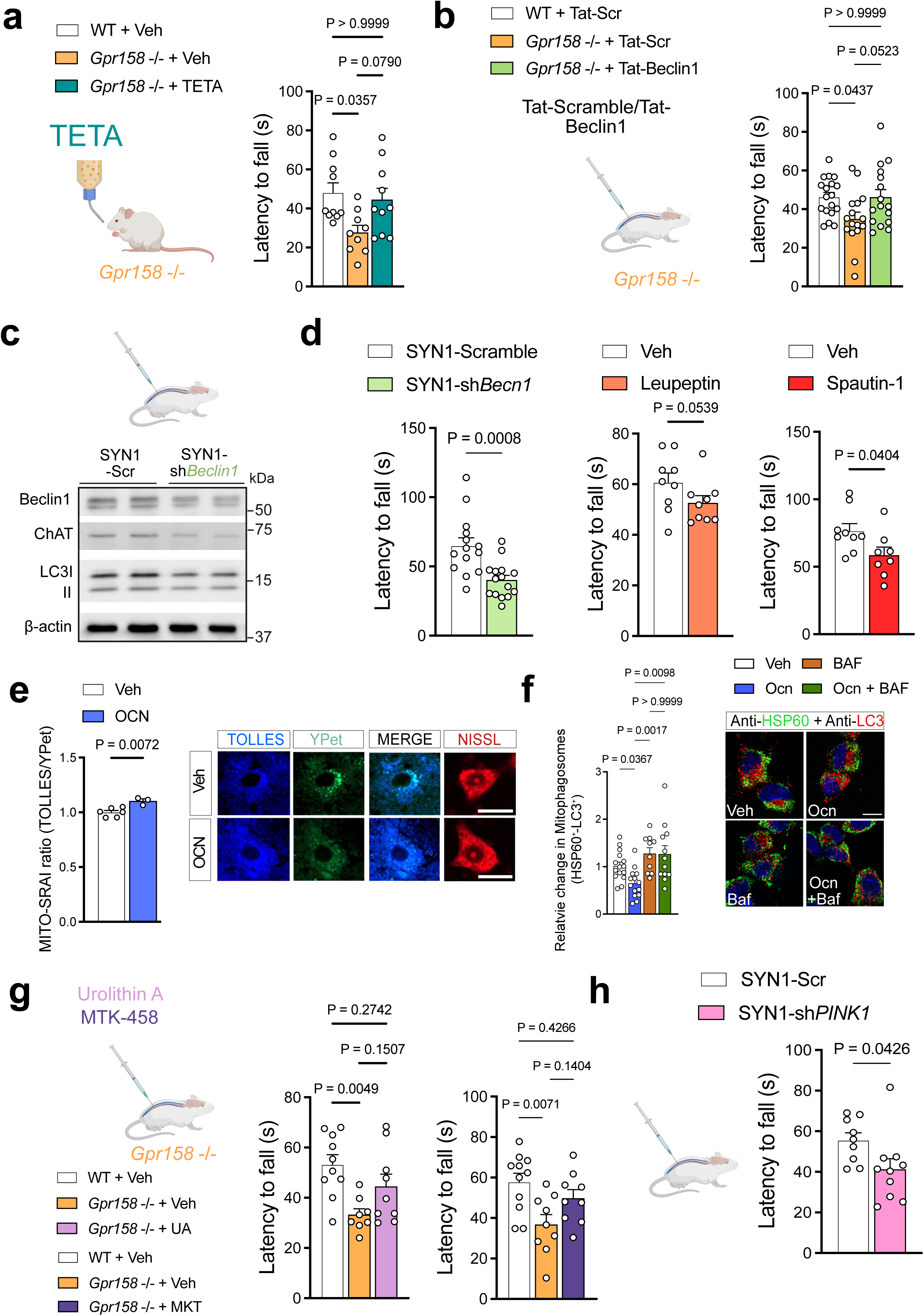
GPR158-dependent mitophagy sustain locomotion functions. **a-** Rotarod performance test in 3-month wild-type (WT) and *Gpr158* -/- male systemically treated during 1 month with TETA. **b-** Rotarod performance test in 3-month wild-type (WT) and *Gpr158*-/- male upon intrathecal injections of TAT-Scramble or TAT-Beclin1. **c-** Western blot of Beclin1, ChAT and LC3b from lumbar spinal cord of 3-month-old mice upon intrathecal injections of AAVrh10-SYN1-Scramble or *shBeclin1.* **d-** Rotarod performance test in 3-month wild-type (WT) upon intrathecal injection of AAVrh10-SYN1-Scramble or *shBeclin1,* or Leupeptin or Spautin-1. **e.-** Representative images and quantification of mito-SRAI ratio (Tolles/YPet) in the spinal cord motoneurons upon vehicle or OCN intrathecal injection. Scale bar = 20 µm. **f.-** Representative images and quantification of NSC34 treated with Ocn and BAF (2 hours) and stained for HSP60 (Green) and LC3b (Red). Colocalization was performed counting HSP60^+^ in LC3b^+^. Scale bar = 10 µm **g.-** Rotarod performance test in 3-month wild-type (WT) and *Gpr158* -/- male upon intrathecal injections of Urolithin-A (UA) or MKT-458 to induce mitophagy. **h-** Rotarod performance test in 3-month wild-type (WT) upon intrathecal injection of AAVrh10-SYN1-Scramble or *shPink1.* Data are expressed as mean ± SEM. The P values were determined by the Kruskal-Wallis test (Dunn’s multiple comparison test) (**a, f**), One-way ANOVA (Tukey’s Dunnett’s multiple comparison test) (**b, g**), Student’s t-test (**d, e**, **h**) or Mann-Whitney test (**d**). Each dot represents an independent mouse (Spinal cord for molecular biology, or behavioral test).

We next asked whether mitophagy process, the selective removal of mitochondria, which was highlighted in our proteomics data. In support of this notion, Ocn-/-, Gpr158 -/- and mice downregulated for Gpr158 in the spinal cord MNs presented lower mRNA levels of the essential mitophagy genes, Pink1 and Parkin2 (**Fig. S5e**). An AAVrh10-SYN1-mito-SRAI reporter was injected into the spinal cord to measure mitophagy flux (Katayama et al., 2020). OCN increased the TOLLES/YPet ratio of the reporter, indicating enhanced levels of mito-autophagolysosomes in spinal MNs (**Fig. 5e**). We also co-treated NSC34 cells with OCN and bafilomycin A1 (BAF) and quantified, the number of LC3 particles colocalising with HSP60 marker as a mitochondrial protein. OCN reduced the number of mito-containing autophagosomes, an effect abolished by BAF, confirming that OCN engages mitophagy (**Fig. 5f**). Conversely, treatment of Gpr158-/-(Li et al., 2025; Ryu et al., 2016)Haga clic o pulse aquí para escribir texto. mice with urolithin A or MTK-458, two mitophagy inducers, rescued rotarod performance, indicating that GPR158-dependent mitophagy influences motor functions (**Fig**. **5g, f**). To confirm the requirement of basal mitophagy in vivo, we downregulated Pink1, a key initiator of mitophagy, specifically in spinal neurons (AAVrh10-SYN1-shPink1) (**Fig**. **S5f**). Mice with shPINK1 intrathecal injection displayed impaired rotarod performance without changes in Open Field locomotion (**Fig. 5h and S5g**). Together, these results establish that OCN signaling maintains motor function through autophagy and, more specifically, mitophagy in spinal MNs.

### OCN–GPR158 regulates mitochondrial activity in spinal motor neurons

Since mitophagy ultimately serves as a mitochondrial quality control, we asked whether OCN could modulate mitochondrial activity. When studying NSC34 MN-like cells using Seahorse extracellular flux analysis, we found that OCN significantly increased basal and maximal oxygen consumption rates (OCR), indicating that OCN enhances mitochondrial respiratory capacity (**Fig. 6a**).

**Figure 6:**
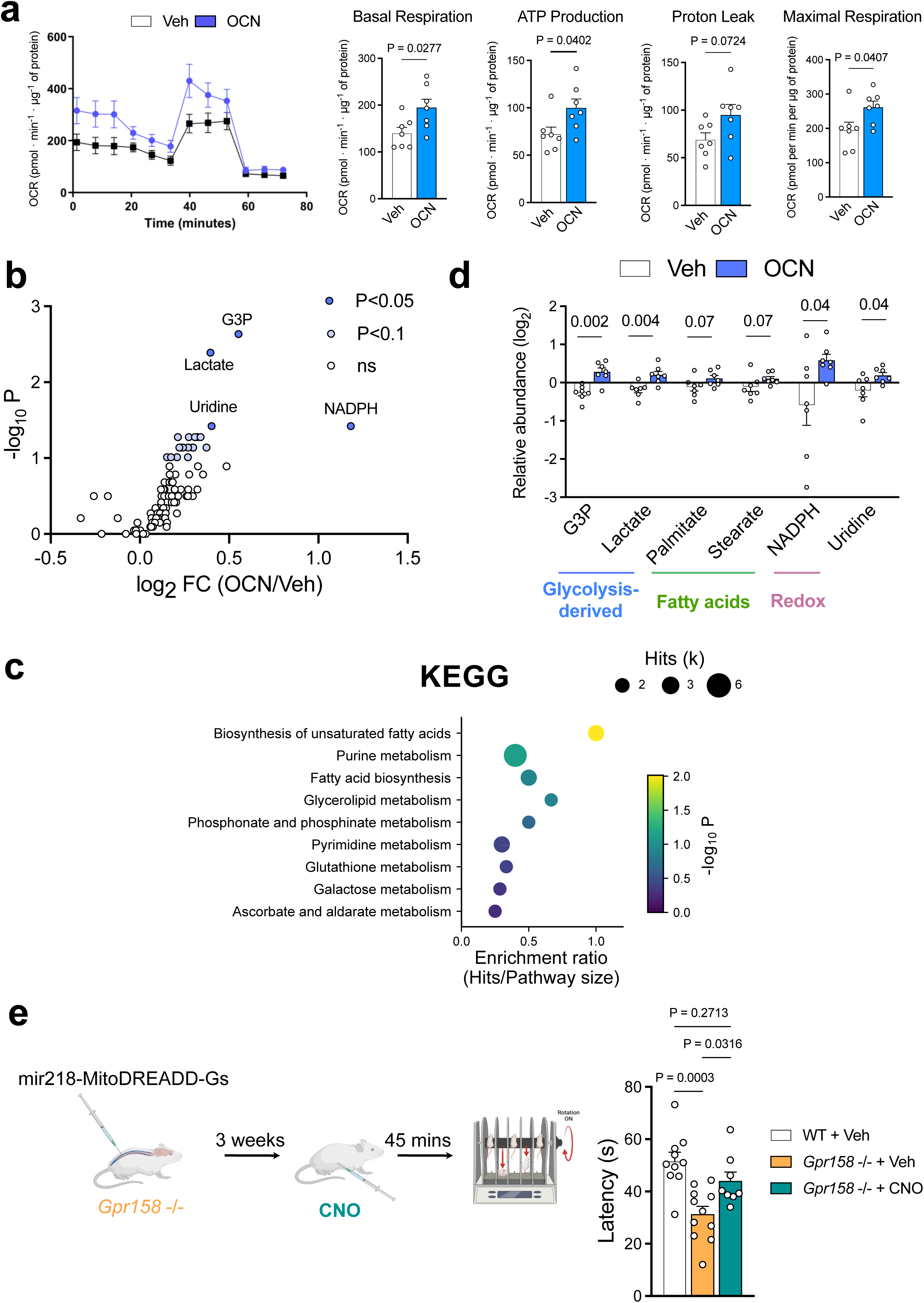
OCN and GPR158 regulate mitochondrial activity. **a.-** Oxygen consumption rate (OCR) assessed by Seahorse in NSC34 treated during 1 hour with OCN. **b.-** Volcano plot of targeted metabolomics from lumbar spinal cord of 3-month-old mice after intrathecal injection of Vehicl or OCN. **c.-** KEGG pathway enrichment analysis of OCN-regulated metabolites. **d.-** Relative abundance of representative metabolites grouped by substrate class (glycolytic: G3P, lactate; fatty acids: palmitate, stearate; redox/nucleoside: NADPH, uridine) in Veh. and OCN groups. **e.-** Rotarod performance in WT and *Gpr158-/-*, upon AAVrh10-mir218-mitoDREADD intrathecal injection, and 45 minutes after Vehicle or CNO intraperitoneal injection. Data are expressed as mean ± SEM. The P values were determined by Student’s t-test (**a**), Mann-Whitney test (**b**, **d**), hypergeometric test (**c**), or one-way ANOVA (Dunnett’s multiple comparison test) (**e**). In (c), pathways with at least two hits are shown. Each dot represents an independent mouse (Spinal cord for molecular biology or behavioral test) an NSC34 independent preparation.

Targeted metabolomics on spinal cord tissue after intrathecal OCN injection revealed that OCN triggers a broad bioenergetic response in the lumbar spinal cord, in which every OCN-responsive metabolite was increased. OCN significantly increased glycerol-3-phosphate and lactate, which feed the glycerophosphate shuttle and the TCA cycle, respectively, as well as the redox cofactor NADPH and the nucleoside uridine. In parallel, the saturated fatty acids palmitate and stearate, together with arachidonate and caprate, tended to increase, pointing to a broader mobilization of lipid substrates. The same trend was observed for several purine intermediates, including adenosine, inosine, hypoxanthine and IMP, and for ribose-5-phosphate and UDP-glucose. KEGG pathway analysis identified biosynthesis of unsaturated fatty acids as the most enriched pathway among the OCN-responsive metabolites, followed by purine metabolism (**Fig. 6b-d and S6a, b**). These data indicate that OCN increases the availability of glycolysis-derived and lipid substrates in the spinal cord, in line with the higher mitochondrial respiration induced by OCN.

To test whether potentiated mitochondrial function is sufficient to recover motor function, we expressed a mitochondria-targeted DREADD (Pagano Zottola et al., 2025) (mitoDREADD-Gs-HA) in spinal MNs of Gpr158-/- mice and activated it chemogenetically, thereby increasing mitochondrial bioenergetics via cAMP. We observed that direct mitochondrial activation rescued rotarod performance, and increased pPKA, indicating DREADD activation (Pagano Zottola et al., 2025) (**Fig. 6e, S6c**). These results indicate that OCN signaling sustains mitochondrial function in spinal MNs, and that potentiating mitochondrial function is sufficient to maintain motor function.

### OCN improves age-related locomotor decline by selective degradation of mitochondria through autophagy

Circulating OCN decrease during aging in all species tested, and restoring OCN improves neuronal functions such as cognition (Khrimian et al., 2017). We therefore asked whether OCN would improve locomotor function through spinal cord MNs. At 16 months of age, mice showed a significant loss of their motor function (**Fig. S7a**), assessed by rotarod and beam bar. Conversely, spinal expression of Gpr158, BDNF, CREB, ChAT, VAChT and Pink1 is reduced in aged mice compared to young (3Mo), combined with lower ChAT levels (**Fig. 7a, b, and S7b**). The autophagy machinery is also reduced with age, as shown by decreased Beclin1 and LC3-II protein levels and increased p62 puncta in spinal cord MNs (**Fig. 7c)**, suggesting that an age-related decline of autophagy exists in these neurons. We therefore asked whether restoring autophagy or OCN could counteract this age-related motor decline. We found that systemic treatment of aged mice (16 months) with TETA improved locomotor performance at rotarod (**Fig. 7d**). OCN chronic administration to aged mice (16 months) significantly improved rotarod and beam bar performance compared to vehicle-treated controls, and restored ChAT levels in the lumbar spinal cord (**Fig. 7e, f S7c**). This indicates that the age-related motor decline can be functionally reversed by OCN.

**Figure 7:**
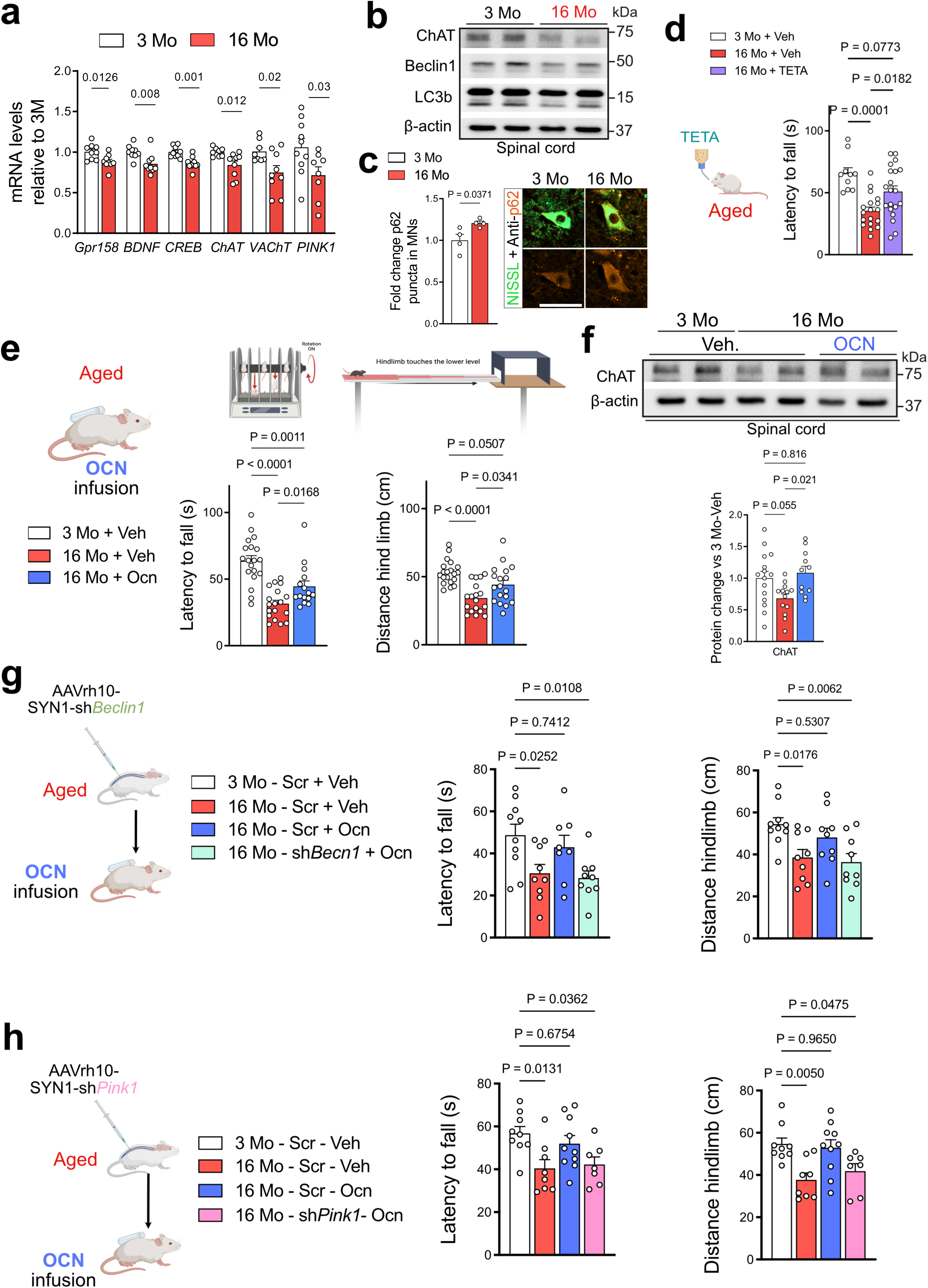
Osteocalcin ameliorates age-related locomotor decline through mitophagy machinery in the spinal cord motoneurons. **a.-** Relative mRNA levels of *Gpr158, BDNF*, *BDNF, ChAT, VAChT, Pink1,* in the lumbar spinal cord from 3- or 16Month-old mice **b.-** Immunoblot for ChAT, Beclin1 and LC3b in the lumbar spinal cord from 3- or 16Month-old mice. **c.-** Spinal cord labelling against p62 (Orange) and FluoroNissl (Green) from 3- or 16Month-old mice, and p62 puncta quantification. Scale bar= 50 µm **d.-** Rotarod performance in 3 and 16Month-old mice upon 1 month of systemic TETA treatment. **e.-** Rotarod performance and beam bar coordination test in 3 and 16Month-old mice upon 1 month of systemic Vehicle or OCN treatment by osmotic pump implantation. **f.-** Representative western blot image and quantification of ChAT in the lumbar spinal cord from 3Mo and 16Mo after the systemic treatment with OCN. **g.-** Rotarod performance and beam bar coordination in 3-month-old and 16-month-old mice that received systemic OCN treatment 3 weeks after the intrathecal injection of intrathecal injection of AAVrh10-SYN1-Scramble or *shBeclin1.* **h.-** Rotarod performance and beam bar coordination in 3-month-old and 16-month-old mice that received systemic OCN treatment 3 weeks after the intrathecal injection of intrathecal injection of AAVrh10-SYN1-Scramble or *shPink1.* Data are expressed as mean ± SEM. The P values were determined by Student’s t-test (**a**, **c**) or Mann-Whitney test (**a**) or one-way ANOVA (Tukey’s, Holm-Šídák’s, Bonferroni’s or Dunnett’s multiple comparison test) (**d, e, f, g, h**). Each dot represents an independent mouse (Spinal cord for molecular biology, or behavioral test).

To determine if this rescue relies on the mechanism identified in young mice, we blocked autophagy or mitophagy specifically in spinal MNs. Intrathecal injection of AAVrh10-Syn1-shBeclin1 or Syn1-shPink1 abolished OCN-induced locomotor improvement in rotarod and beam coordination test (**Fig. 7g, h, S7d, e**), demonstrating that OCN’s anti-aging action requires autophagy and mitophagy machinery. Together, these data define a bone-to-spinal cord endocrine axis in which OCN signaling through GPR158 sustains motoneuron mitochondrial quality and locomotor performance across lifespan.

## Discussion

In this study, we identify an endocrine pathway through which the skeleton regulates motor function trough spinal cord. Our data establish that bone, by producing OCN, is an active endocrine component of the locomotor system and suggest that bone-to-spinal cord communication contributes to locomotor resilience during aging.

When we examined the full landscape of hormonal receptors in spinal cholinergic neurons, alpha MNs emerged as the spinal population most enriched in, and most dependent on, GPCR signaling. This was not limited to GPR158, suggesting that alpha MNs are, as a class, unusually dependent on GPCR-mediated signaling. This broad enrichment of GPCR signaling suggested that circulating ligands could represent an underappreciated layer of regulation of spinal motor function.

The focus on OCN was motivated by its unique biology as a bone-specific hormone released in response to skeletal mechanical loading (Hiam et al., 2021; Mera, Laue, Ferron, et al., 2016). The muscle is also a direct target of OCN (Mera, Laue, Ferron, et al., 2016; Mera, Laue, Wei, et al., 2016), and the locomotor system depends on the coordinated function of all three: bone, which produces OCN; the spinal cord, whose MNs we now show respond to it; and muscle, whose contraction depends on the bone and is regulated by OCN. OCN may therefore provide an endocrine link between skeletal loading, systemic energy demands, and the functional state of the locomotor system. The age-related decline in circulating OCN (Mera, Laue, Ferron, et al., 2016; Smith et al., 2020) could contribute to motor deterioration not only through effects on bone and muscle mass, but also through loss of endocrine support to spinal MNs, potentially compromising their bioenergetic capacity (Rygiel et al., 2014).

Our genetic and cell-type-specific approaches point to GPR158 as the primary mediator of OCN effects in spinal MNs. GPR158 is selectively enriched in ChAT-positive MNs, its loss phenocopies OCN deficiency, and the molecular effects of OCN are abolished in Gpr158-deficient mice. These findings identify GPR158 as a key receptor through which circulating OCN is translated into a cell-autonomous response in spinal MNs. However, we cannot exclude a potential contribution from GPR37, a recently proposed OCN receptor expressed in spinal oligodendrocytes and some neurons (Bian et al., 2024; Liu et al., 2026; Qian et al., 2022). Whether OCN-GPR37 signaling in these populations contributes to motor function remains an open question that will require cell-type-specific models.

A key mechanistic question raised by our data is whether the effects of OCN on mitophagy and mitochondrial function represent a single integrated program or two parallel responses. Blocking mitophagy machinery abolishes the beneficial effects of OCN, supporting an essential role for mitophagy in the OCN response. Yet mitoDREADD, which activates mitochondria directly without engaging autophagy, is sufficient to rescue the *Gpr158-/-* motor phenotype, indicating that enhanced mitochondrial activity can bypass the functional consequences of GPR158 loss. We therefore propose that OCN-GPR158 may couple two complementary aspects of mitochondrial homeostasis: selective removal of dysfunctional mitochondria through mitophagy and enhancement of mitochondrial bioenergetic capacity through a parallel signal, potentially involving Ca^2+^ signaling or SIRT1-dependent mechanisms, as observed in the hippocampus (Rivagorda et al., 2025; Romeo-Guitart et al., 2026). These processes are not necessarily redundant: mitophagy clears damaged mitochondria, whereas metabolic activation may potentiate the functional capacity of the remaining mitochondrial pool. Together, these mechanisms could sustain the energetic demands of MNs and consequently motor function, a response conceptually analogous to the mitochondrial adaptations induced by exercise in skeletal muscle (Hood et al., 2015; Laker et al., 2017).

The exceptional axonal length of MNs makes mitochondrial quality control particularly consequential in this cell type. Alpha MNs have axons exceeding one meter in humans required to innervate distal muscles (Dadon-Nachum et al., 2011; Kanning et al., 2010), and mitochondria must be transported from the soma to terminals where energy demand is highest. Dysfunctional mitochondria that are not cleared accumulate along the axon and at neuromuscular junctions, impairing transmission. Thus, disruption of the OCN-GPR158 pathway may be particularly detrimental to MNs because these cells combine exceptionally long axons with sustained energetic demands. Consistent with this model, spinal cord metabolomics shows that OCN increases several metabolites linked to mitochondrial energy metabolism at once: G3P and lactate, the redox cofactor NADPH and, saturated fatty acids, which provide oxidative substrates and support redox balance (Herrero-Mendez et al., 2009; Magistretti & Allaman, 2015). This coordinated metabolic signature is consistent with the increase in mitochondrial function induced by OCN-GPR158, rather than reflecting a change in a single metabolic pathway. Because OCN is released during physical activity, this metabolic response further supports a model in which OCN helps match spinal MN bioenergetic capacity to physiological demand.

Our findings are consistent with the emerging notion that the benefits of physical exercise on motor aging may be mediated in part through OCN. Exercise increases circulating OCN through mechanical loading of bone (Hiam et al., 2021; Horowitz et al., 2020; Mera, Laue, Ferron, et al., 2016), and OCN in turn maintains spinal MN function through mitochondrial quality control. This positions bone not simply as a structural component of the locomotor system, but as an endocrine organ capable of communicating the physiological consequences of physical activity to the nervous system.

Systemic OCN restores motor performance in aged mice. The pathway therefore remains responsive with aging, and reduced endocrine input may contribute to age-related motor decline. The requirement of mitophagy machinery in MNs for OCN to exert its protective effects during aging aligns with the importance of mitochondrial quality control and mitophagy for neuronal aging and neurodegeneration in other systems (Grimm et al., 2026; Pan et al., 2026; Ryu et al., 2016; Wood-Kaczmar et al., 2008). Indeed, exercise-induced mitochondrial turnover in skeletal muscle is among the most reproducible interventions against age-related functional decline (Hood et al., 2019; Laker et al., 2017). Together, these findings suggest that restoring an endogenous endocrine signal can reinforce mitochondrial quality-control mechanisms that become insufficient during aging, providing a potentially reversible component of motor decline.

In summary, we demonstrate that the skeleton regulates locomotion through OCN signaling in spinal motor neurons, thereby maintaining mitochondrial function. This axis declines with age in parallel with circulating OCN (Mera, Laue, Ferron, et al., 2016) and restore OCN is sufficient to reverse age-related motor decline. Our findings establish a bone-spinal cord communication pathway that links skeletal physiology to mitochondrial homeostasis in spinal MNs and identify endocrine regulation as an additional layer of control over motor aging. More broadly, this work reveals that peripheral endocrine signals can directly shape the resilience of spinal motor circuits, providing a new framework for understanding how systemic physiology contributes to the maintenance of motor function throughout life.

## Material and Methods

### Animals

All the experiments were performed on C57BL/6J WT male mice (Janvier Laboratory, Le Genest St Isle, France), except the ones using *Bgp*-/- (129Sv-se background as described (Oury et al., 2013) or *Gpr158*-/- (C57BL/6J background as used in (Khrimian et al., 2017)), that were performed in male and in female. Mice were randomly distributed in cages independently of their genotype. We used littermates as controls for all experiments with the Bgp or Gpr158 knock-out strains. Mice were 3 or 16 months of age at the start of experiments. Upon arrival, mice were housed at least 2 weeks before any intrathecal injection, osmotic pump implantation, or behavioral and molecular testing, under a 12-hour light/dark cycle with ad libitum access to water and food. All the behavioral experiments were performed in the morning (between 9am and 1pm). Animals were randomly assigned to different groups before starting the experiments. All behavioral experiments were performed in accordance with the European Communities for Experimental animal use (2010/63/EU) and local ethical committee review procedures and protocols (APAFIS).

### Intrathecal Injection

Intrathecal delivery of drugs, OCN, and viral particles was carried out at the lumbar level in anaesthetized mice using a Hamilton syringe fitted with a 33-gauge needle, as described previously(Romeo-Guitart et al., 2018, 2019). After locating the vertebrae through an incision in the lumbar region, 10 µl were injected slowly into the CSF between the L3 and L4 vertebrae. Correct positioning within the intrathecal space was verified by a tail flick response. The needle was left in place for one minute after injection, and the muscle and skin were then sutured.

### Osmotic pumps implantation

For the chronic treatment with OCN, we filled Alzet micro-osmotic pumps with either vehicle (0.9% NaCl) or uncarboxylated osteocalcin (ucOCN), delivered at 30 ng h⁻¹. Mice were anaesthetised with isoflurane; we performed an incision on the dorsal region and implanted the pumps subcutaneously as described (Rivagorda et al., 2025). Incisions were closed with sutures, and animals were placed in a heated recovery chamber and monitored until ambulatory activity was fully recovered. Behavioural testing began two weeks after implantation.

### GPCR analysis from single-nucleus RNA sequencing

We used available single-nucleus RNA sequencing data of the adult mouse spinal cord from the Spinal Cord Atlas (spinalcordatlas.org) (Alkaslasi et al., 2021; Yadav et al., 2023). To dissect both cell-type-specific vulnerability and baseline network architecture, we combined a targeted analysis of non-redundant GPCR hubs from specific cell-type-specific genes with a broader examination of the complete gene expression profile.

By analytically focusing on cell-type-specific genes, transcriptomic data was filtered for significance (p < 0.05), and duplicate gene entries across clusters were resolved using the maximum average log-fold change. Protein-protein interaction (PPI) interactomes between specific neuron types and G Protein-Coupled Receptors (GPCRs) were mapped using the STRING database with a high-confidence interaction score threshold of 0.7, categorized according to KEGG classification (mmu04030). To assess the structural significance and biological vulnerability of each motor neuron subtype, a systematic node-deletion sensitivity analysis was performed. The degree change was calculated by comparing the mean degree centrality of the baseline graph against the graph resulting from the iterative removal of each subtype-specific node. Global centrality shifts following *in silico* gene deletions were analyzed using box plots to capture the heavy-tailed distributions of high-impact genes. Pairwise differences in network vulnerability across neuronal populations were evaluated using the Mann-Whitney U test. To characterize the molecular identity of structurally significant receptors, GPCRs within the alpha motor neuron interactome were ranked by degree change and partitioned into five equal-sized bins (ranging from Bin 1 for most critical impactors to Bin 5 for least impactful nodes). The percentage distribution of GPCR functional families across these bins was evaluated using stacked bar charts.

To capture the overall, baseline physiological wiring of the network beyond distinct subpopulation markers, the complete gene expression profile was analyzed using processed single-cell spinal motor neuron sequencing data (.h5ad matrices with cluster IDs and annotations). Mouse GPCR symbols, hierarchical levels, and family annotations were retrieved via the KEGG REST API into a structured Data Frame and cross-referenced against the genes present in the dataset. To standardize expression profiles across diverse cell populations, row-wise Z-scores were calculated for each GPCR to reflect relative enrichment rather than absolute transcript abundance. Hierarchical clustering of these Z-scored profiles identified a distinct transcriptional module (Cluster 6, which includes *Gpr158*) characterized by marked positive Z-scores and selective upregulation in motor neuron populations. To evaluate how target cluster expression varies across lineages, the mean expression per cell for cluster genes was computed, and pairwise independent t-tests were performed across all cell types. Compact letter displays (significance threshold of p < 0.001) and -log(p-value) heatmaps were generated to map statistical separation between high-expressing neuronal groups and lower-expressing glial and endothelial populations. A dedicated evaluation of *Gpr158* expression was similarly performed to assess its mean counts and distribution across all cell types.

### NSC34 cell culture, transfection, autophagy assessment and Immunofluorescence

NSC34 cells were seeded at a density of 1.5 × 10⁵ cells/ml in 24-well plates. For transfection, 24 hours following seeding, cells were transfected with 500ng/well of the construct expressing Scr or *shGpr158* using Lipofectamine 2000 transfection reagent (Thermo Fisher Scientific), according to the manufacturer’s protocol, with minor adaptations. Cells were incubated with the transfection mixture for 6 hours, and then the media was changed to differentiation medium consisting of DMEM/F12 (1:1) supplemented with 1% FBS, 0.5% penicillin/streptomycin, and 10 µM retinoic acid (RA). Western blot and Immunofluorescence experiments were performed 4 days after transfection. NSC34 were treated with OCN at a final concentration of 10ng/ml and Bafilomycin (BAF) at 10nM final concentration. Cells were harvested at 4 hours for Western blot and at 1 hour for immunofluorescence to assess mitophagy.

### Spinal cord proteomics

S-Trap^TM^ micro spin column (Protifi, Hutington, USA) digestion was performed on 50µg of spinal cord cell lysates according to manufacturer’s instructions. Briefly, samples were reduced with 20mM TCEP (Tris(2-carboxyethyl) phosphine hydrochloride) and alkylated with 50mM CAA (chloracetamide) for 5min at 95°C. Aqueous phosphoric acid was then added to a final concentration of 2.5% following by the addition of S-Trap binding buffer (90% aqueous methanol, 100mM TEAB, pH7.1). Mixtures were then loaded on S-Trap columns. Six washes were performed for thorough SDS elimination. Samples were digested with 2.5µg of trypsin (Promega) at 47°C for 1h and 30 minutes. After elution, peptides were vacuum dried and resuspended in 50µL of 2% ACN, 0.1% formic acid in HPLC-grade water prior to MS analysis.

400 ng of each sample was injected on a nanoElute (Bruker Daltonics, Germany) HPLC (high-performance liquid chromatography) system coupled to a timsTOF Pro (Bruker Daltonics, Germany) mass spectrometer. HPLC separation (Solvent A: 0.1% formic acid in water; Solvent B: 0.1% formic acid in acetonitrile) was carried out at 250nL/min using a packed emitter column (C18, 25 cm×75μm 1.6μm) (Ion Optics, Australia) using a 70min gradient elution (2 to 13% solvent B during 42min; 13 to 20% during 23min; 20% to 30% during 5min; 30% to 85% for 5min and finally 85% for 5min to wash the column).

Mass-spectrometric data were acquired using the parallel accumulation serial fragmentation (PASEF) acquisition method in DIA mode. The measurements were carried out over the m/z range from 100 to 1700 Th. The range of ion mobility values was from 0.6 to 1.6 V s/cm2(1/k0). diaPASEF settings were: mass range from 400 to 1200 m/z, mobility ranges from 0.60 to 1.43 V s/cm2 1/k0, cycle time estimate of 1.79 s, 16 PASEF scans x 2 steps per PASEF scans (32 windows).

Data analysis was performed using DIA-NN software (version 1.8.1) (Demichev et al., 2020). A search against the UniProtKB/Swiss-Prot Mus musculus database (release March 2022, 17515 entries) was performed using a library free workflow. For this purpose, “FASTA digest for library free search/library generation” and “Deep learning spectra, RTs and IMs prediction” options were checked for precursor ion generation. 2 trypsin missed cleavage was allowed and the maximum variable modification was set to 3. Carbamidomethylation (Cys) was set as the fixed modification, whereas protein N-terminal methionine excision, methionine oxidation and N-terminal acetylation were set as variable modifications. The peptide length range was set to 7–30 amino acids, precursor charge range 2–4, precursor m/z range 300–1800, and fragment ion m/z range 200–1800. To search the parent mass and fragment ions, accuracy was set to 10 ppm manually. The false discovery rates (FDRs) at the protein and peptide level were set to 1%. Match between runs was allowed. For the quantification strategy, Robust LC (high precision) was used as advised in the software documentation, whereas default settings were kept for the other algorithm parameters.

Statistical and bioinformatic analysis were performed with Perseus software (version 1.6.15.0) (Tyanova et al., 2016),freely available at www.perseus-framework.org. All protein intensities were log2-transformed to perform statistics. For statistical comparison, we set three groups: WT+Veh, WT+OCN, and *Gpr158* -/-, each containing 5 biological replicates. Proteins were retained for analysis only if they were quantified in at least 4 of the 5 samples in at least one group. Next, the data were imputed to fill missing data points by creating a Gaussian distribution of random numbers with a standard deviation of 33% relative to the standard deviation of the measured values and 1.8 standard deviation downshift of the mean to simulate the distribution of low signal values. We performed one-way ANOVA with FDR<0.05 and S0 parameter at 0.1. Hierarchical clustering of proteins that were significant was performed in Perseus after z-score normalization of the data, using Euclidean distances.

#### Differential expression analysis

To quantify differential expression across the experimental groups (WT, WT+OCN, and *Gpr158*-/-), we used an adapted version of the DESeq2 package in R/Bioconductor. The dataset was normalized using log and normalized log transformations, excluding any data entries in the WT and WT+OCN groups where reads were zero across more than 75% of the samples within an experimental group. Principal component analysis (PCA) was performed on the DESeq2 count-read object using the prcomp function in R to identify outliers, reduce dimensionality, and estimate the proportion of variance explained by orthogonal eigenvectors derived from Euclidean distance metrics. The underlying DESeq2 methodology follows (Love et al., 2014)

For each protein, a logarithmic generalized linear model (GLM) was fitted to evaluate baseline expression strength, maximum-likelihood estimates (MLEs) for log2 fold changes (LFC) between groups (e.g., WT vs. WT+OCN), and gene-wise dispersion estimates via maximum likelihood. A plot of data count dispersion against average expression revealed the distribution of these estimates, which were then refined using an empirical Bayes approach to determine degrees of freedom and fit accuracy. Standard GLMs were run to calculate LFC MLEs, followed by fitting a zero-centered normal distribution across all protein MLEs. Count data were transformed using either regularized logarithmic transformation (rlog) which controls for differences in sequencing depth or variance-stabilizing transformation (vst) for overdispersed data. Outliers post-GLM fitting were flagged using Cook’s distance, measuring the coefficient vector shift upon single-sample exclusion. Statistical significance of the LFC was evaluated using a Wald test, where shrunken LFC estimates were divided by their standard error to generate z-statistics compared against a standard Gaussian distribution. The resulting p-values were adjusted for multiple testing using the Benjamini-Hochberg procedure, with significantly up-regulated and down-regulated proteins (FDR = 10%) highlighted in purple and blue, respectively.

#### Weighted correlation gene network analysis

Weighted Gene Co-expression Network Analysis (WGCNA) was performed to identify functional co-expression networks and module patterns across protein/gene expression profiles. When two genes/proteins show similar expression patterns across multiple samples such as, both increasing or decreasing together, they are said to be co-expressed. This co-expression often indicates functional relationships: genes working together in the same pathway, responding to the same regulatory signals, or contributing to the same biological process. By representing individual genes or proteins as nodes and their pairwise correlation strength as edges, WGCNA preserves continuous co-expression relationships to infer biological pathways, shared regulatory mechanisms, and co-functional networks.

Network construction evaluated expression profiles across all WT, WT-OCN and *Gpr158*-/- samples, pooling across all experimental ages. Features with missing values in over 50% of replicates within a sample-age group were removed. A pairwise correlation coefficient matrix was generated and transformed into a weighted adjacency matrix using a soft-thresholding power (β) of 10. This soft thresholding suppressed weak correlations while highlighting strong connections, ensuring the network achieved a scale-free topology characteristic of biological systems. To reduce noise and account for shared neighborhood connections, the adjacency matrix was converted into a Topological Overlap Measure (TOM).

Using TOM dissimilarity (1 – TOM) as a distance metric, hierarchical clustering was conducted to partition genes/proteins into co-expression modules. Using a dynamic tree-cut threshold of 0.55, the algorithm automatically resolved 8 distinct modules containing genes with high topological overlap. For each identified module, a module eigengene (ME), defined as the first principal component of the module’s expression matrix, was computed to summarize the overall expression trajectory. MEs provide representative metrics that can be correlated with external phenotypic traits, offering a robust framework for hypothesis generation and mechanistic discovery even when experimental data are sparse or confounded. The methodology has been adapted previously in other studies and addressed key biological network components effectively (Ramos-Brossier et al., 2023; Rivagorda et al., 2025). Based on the global Z-scores across the modules/clusters, the clusters were categorized into 4 broad clusters based on their mean change of the Z-scores (Cat 1-4).

#### Protein-protein interaction network

To model the functional interactions and cross-talk between biological pathways, a pathway-level graph was constructed using protein-protein interaction (PPI) data as a baseline scaffolding. The reference PPI network was retrieved from the STRING database(Szklarczyk et al., 2023), applying a high-confidence threshold (combined score 0.7). Biological pathways were subsequently mapped to the network nodes by cross-referencing constituent genes with their respective functional pathways. A subnetwork comprising all genes associated with the target pathways was extracted for downstream graph projection. The gene-level PPI network was abstracted into a weighted, undirected pathway cross-talk graph, where the set of vertices represents individual pathways. Pathway-pathway edges were established based on the presence of shared PPIs. These edges were weighted by the total frequency of inter-pathway protein interactions to represent cross-talk density. Ultimately, normalized weighted degree centrality was computed for each node to highlight the key biological pathways governing the network structure.

#### Ontology analysis

To characterize the biological functions of genes identified within the experimental group-dependent clusters (WT, WT+OCN, and *Gpr158*-/-), functional enrichment analysis was performed using an over-representation approach based on the hypergeometric distribution. Candidate gene lists were evaluated against three curated annotation databases: the Kyoto Encyclopedia of Genes and Genomes (KEGG), Reactome, WikiPathways, CORUM and Gene Ontology (GO). GO analyses were systematically divided across the primary domain category of the Biological Process (BP). All enrichment tests were configured for *Mus musculus*. To control the false discovery rate (FDR) arising from multiple hypothesis testing, p-values were adjusted using the Benjamini-Hochberg procedure. Pathways and terms satisfying both an adjusted p-value (ρ_adj_) threshold of <0.05 and a q-value threshold of <.05 were considered statistically significant. The broad processes were grouped into categories of pathways (glucose metabolism, immune and inflammation, oxidative phosphorylation, TCA cycle, MAPK signaling, fatty acid metabolism, Mitochondria related processes and nucleic acid metabolism) based on a custom NLP algorithm designed to classify processes as follows:

- Glucose Metabolism: Encompasses pathways controlling carbohydrate uptake, breakdown, and interconversion, including glycolysis, gluconeogenesis, hexose/monosaccharide transport, and pyruvate metabolism.
- Immune and Inflammation: Encompasses innate and adaptive immune responses, cytokine and chemokine signaling pathways, inflammatory cascades (e.g., TNF and NFkB signaling), and lymphocyte-mediated pathways (e.g., T-cell and B-cell receptor signaling).
- Oxidative Phosphorylation: Covers respiratory chain complexes, electron transport chain cascades, ATP synthase activity, and cellular oxygen sensing.
- TCA Cycle: Encompasses enzymatic steps and metabolic intermediates of the tricarboxylic acid (citric acid/Krebs) cycle, including succinate, fumarate, and malate processing.
- MAPK Signaling: Includes Mitogen-Activated Protein Kinase signaling cascades, as well as crosstalk through p53, RAF, JNK, ERK, and p38 pathways.
- Fatty Acid Metabolism: Encompasses fatty acid oxidation (beta-oxidation), lipogenesis, lipid transport, and broader lipid/phospholipid metabolic processes.
- Mitochondria Related Processes: Captures mitochondrial structural integrity, cristae organization, mitochondrial membrane dynamics, and respiratory complex assembly.
- Nucleic Acid Metabolism: Encompasses purine and pyrimidine biosynthesis and salvage, nucleoside/nucleotide interconversion, and nucleobase metabolic pathways.

Pathways matching keywords in multiple superclasses were assigned to all matching process categories.

#### Pathway level centrality measurements

The biological connectivity was modeled as two distinct network levels using the *networkx* graph theory library. First, the Global Network (*G*_total_), denoting an undirected co-membership graph containing all unique genes across assigned pathways as nodes. An edge (*g_i_, g_j_*) was established between any two genes if they co-occurred in at least one enriched pathway. For the second layer, local subnetworks (*G*_proc_) consisted of subgraphs constructed independently for each of the eight process superclasses using only the genes mapped to pathways within that respective superclass. To identify critical driver genes at both system-wide and process-specific scales, three primary centrality metrics were evaluated on both *G*_total_ and *G*_proc_:

Degree Centrality (*G_D_*): Measures the direct functional connectedness of a gene *v* based on its degree *k_v_* relative to total reachable nodes *N* – 1:

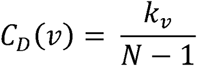

Betweenness Centrality (*C_B_*): Quantifies the degree to which a gene acts as an information bridge along shortest topological paths (*σ_st_*) passing through node *v*:

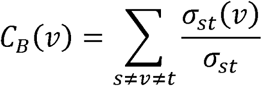

Eigenvector Centrality (*C_E_*): Measures influence by weighing a node’s connections according to the centrality of its neighbors, solved via the principal eigenvector of adjacency matrix *A*:

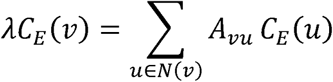

Because individual centrality metrics operate on different mathematical scales, each metric vector *X* ∇ {*C_D_*, *C_B_*, *C_E_*} was mapped to a normalized 0 – 1 domain via Min-Max scaling:

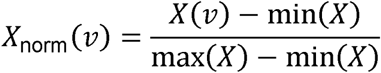

Finally, an integrated centrality score was calculated for both global and local contexts by taking the arithmetic mean of the three scaled metrics:

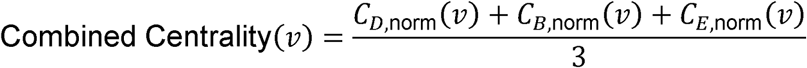

Genes within each process superclass were ranked by local combined centrality, and the top 10 central genes per functional group were selected. The overarching network layout was constructed by taking the induced subgraph (*G*_top_) of these selected genes from the global network *G*_total_. Nodes were assigned discrete categorical colors corresponding to their primary functional superclass, while the size indicated each node’s combined global centrality score to highlight connected hubs. The graphs were generated using *matplotlib* function in Python.

### Spinal cord metabolomic analysis

Tissues were weighted (about 20mg) and transferred to 2 mL-homogenizer tubes with ceramic beads (Hard Tissue Homogenizing CK28, 2.8 mm zirconium oxide beads; Precellys, Bertin Technologies), containing 500 µL of ice-cold extraction mixture (metOH/water, 9/1, -20°C, with a cocktail of internal standards). The samples were homogenized in Precellys 24 tissue homogenize (3 cycles of 20 s/ 5000 rpm), followed by centrifugation (15 min at 10000 g, 4°C), and collection of the supernatants. Supernatants were then collected and divided in several fractions. 1500 μL of supernatant were transferred to glass vials and evaporated to be used for GC-MS analysis. Then, 50 μL of methoxyamine (20 mg/ml in pyridine) was added on dried extracts that were stored at room temperature in the dark, overnight. The day after, 80 μL of N-Methyl-N-trimethylsilyl-trifluoroacetamide were added, and the final derivatization was made at 40°C during 30 minutes. Samples were then transferred into vials and directly injected into the GC-MS system. Another 150 µL of supernatant was transferred into glass vials and evaporated. Dried extracts were spiked with 100 µL of Milli-Q water and directly injected into the LC-MS system for nucleotides, co-factors and related metabolites analysis (Suissa et al., 2026)

#### Widely-targeted analysis of intracellular metabolites (GC-MS)

Widely-targeted analysis of intracellular metabolites was performed using GC-MS/MS method on a 7890B gas chromatography (Agilent Technologies, Waldbronn, Germany) coupled to a triple quadrupole 7000C (Agilent Technologies, Waldbronn, Germany) equipped with a high sensitivity electronic impact source operating in positive mode as previously described (Viltard et al., 2019). Pseudo-targeted analysis of intracellular metabolites (including fatty acids) was performed with a Dionex Ultimate 3000 UHPLC system (Thermo Scientific) coupled to a Q-Exactive (Thermo Scientific) equipped with an electrospray source operating in both positive and negative mode and full scan mode from 100 to 1200 m/z. The QExactive parameters were: sheath gas flow rate 55 a.u., auxiliary gas flow rate 15 a.u., spray voltage 3.3 kV, capillary temperature 300 °C, S-Lens RF level 55 V. The mass spectrometer was calibrated with sodium acetate solution dedicated to low mass calibration (Viltard et al., 2019).

#### Targeted analysis of nucleotides and cofactors by ion pairing ultra-high performance liquid chromatography (UHPLC) coupled to a Triple Quadrupole (QQQ) mass spectrometer

Targeted analysis was performed on a RRLC 1290 system (Agilent Technologies, Waldbronn, Germany) coupled to a Triple Quadrupole 6470 (Agilent Technologies) equipped with an electrospray source.10 μL of sample were injected on a Column Zorbax Eclipse XDB-C18 (100 mm x 2.1 mm particle size 1.8 µm) from Agilent technologies. Gradient mobile phase consisted of water with 2mM of dibutylamine acetate concentrate (DBAA) (A) and acetonitrile (B). Flow rate was set to 0.4 mL/min, and gradient as follow: initial condition was 90% phase A and 20% phase B, maintained during 3 min. Molecules were then eluted using a gradient from 10% to 95% phase B over 1 min. The column was washed using 95% mobile phase B for 2 minutes and equilibrated using 10% mobile phase B for 1 min. Scan mode used was the MRM for biological samples. Peak detection and integration of the analytes were performed using the Agilent Mass Hunter quantitative software (B.10.1) (Solier et al., 2023).

#### Targeted analysis of sugar phosphates by ion pairing ultra high-performance liquid chromatography (UHPLC) coupled to a Triple Quadrupole (QQQ) mass spectrometer

Targeted analysis is performed on a RRLC 1260 system (Agilent Technologies, Waldbronn, Germany) coupled to a Triple Quadrupole 6410 (Agilent Technologies) equipped with an electrospray source operating in negative mode. The gas temperature is set to 325°C with a gas flow of 12 L/min. The capillary voltage is set to 4.5 kV. 10 μL of sample are injected on a Column Poroshell 120 phenyl-hexyl (100 mm x 2.1 mm particle size 2.7 µm) from Agilent technologies, protected by a guard column SB-C18 (5 mm × 2.1 mm particle size 1.8 μm) and heated at 40°C by a pelletier oven. The gradient mobile phase consists of 10 mM tributylamine (TBA) and 10 mM acetic acid (AA) in water (phase A) and in methanol (phase B). The flow rate is set to 0.2 mL/min, and gradient as follow: initial condition is 98% phase A and 2% phase B, maintained during 1 min. Molecules are then eluted using a gradient from 2% to 70% B over 10 min. The column is washed using 95% B for 2 minutes and equilibrated using 2% mobile phase B for 4 min. The scan mode used is the MRM for biological samples. Peak detection and integration of analytes are performed using Agilent Mass Hunter quantitative software (B.07.01).

All targeted treated data were merged and cleaned with a dedicated R package (version 4.0) (https://github.com/kroemerlab/GRMeta). Metabolites were compared between groups with a two-sided exact Mann-Whitney U test (p<0.01), without correction for multiple comparisons. Downstream analyses were performed on the resulting normalized matrix: principal component analysis was computed on the Z-scored metabolite values; for the heatmap, abundances were converted to row-wise Z-scores; and KEGG pathway over-representation analysis was performed on the metabolites with P < 0.1 using a hypergeometric test, with the measured metabolites as background.

### RNAscope for In situ Hybridization

*Gpr158, ChAT, Iba1* and *Gfap* mRNA in-situ hybridization on mouse spinal cord sections was performed with RNAscope Multiplex Fluorescent assay and ACD HybEZ™ II Hybridization System (Biotechne). Briefly, wild-type and *Gpr158*-/- mice were transcardially perfused with cold PBS followed by cold 4% PFA, lumbar segment of spinal cord was harvested, post-fixed 2 h in 4% PFA, cryoprotected in 30% sucrose at 4 °C, embedded in cryo-embedding medium (OCT), and frozen. 12 μm transversal spinal cord sections were processed according to the manufacturer’s instructions. RNAscope probes Mm-Gpr158-C1 was visualized with Opal™ 520, while Mm-ChAT-C2, Mm-Iba1-C2, Mm-Gfap-C2 were visualized with Opal™ 540 Reagents. Slides were counterstained with DAPI and mounted with Fluoromount G. Image acquisition was performed by using a Zeiss Spinning Disk fluorescence microscope (Zen software, Zeiss).

### Western blot analysis

Mouse spinal cord lumbar region or NSC34 cell culture were harvested, lysed in RIPA lysis buffer (25mM Tris HCl, pH 7.6, 150 mM NaCl, 1% NP40, 1% Na deoxycholate, 0.1% SDS and protease and phosphatase inhibitors) and the protein concentration was analyzed in the supernatant by BCA assay. The lysates were loaded on polyacrylamide gel and transferred onto a PVDF membrane. The blots were blocked in Tris-buffered saline with Tween (TBST)-6% milk and incubated with either: mouse anti-β-actin (1:5000, A-2228: Sigma), mouse anti-α-Tubulin (1:5000, T9026, Sigma), mouse anti-GAPDH (1:5000, MAB374: Merck), rabbit anti-GPR158 (1:1000; G307, Assay Biotechnology) mouse anti-Beclin1 (1:1000, 612113: BD Transduction Laboratories, only the top band was quantified), rabbit anti-LC3B (1:10000, L7543: Sigma), rabbit anti-ChAT (1:1000, PA5-29653, Invitrogen), rabbit anti-pPKA (1:1000, 4781, Cell signaling), rabbit anti PKA 1:1000, 4782, Cell signaling). Horseradish peroxidase–conjugated secondary antibodies (anti-mouse or anti-rabbit, HRP-linked antibody, Cell Signaling) were revealed using an ECL kit Clarity Western ECL Substrate (BioRad) or SuperSignal™ West Atto Ultimate Sensitivity Substrate (Thermofisher), for protein detection with the aid of a Bio-Rad ChemiDoc Touch Imaging System version 1.0 and analyzed with BioRad Image Lab software (Version 5.2). β-actin or α-Tubulin were used for normalization. Samples were prepared by one investigator, and the western blot and its analysis was performed by a second investigator blinded to the experimental groups.

### cFOS+ spinal motoneurons counting

Wild-type and *Gpr158*-/- mice were first deeply anesthetized with a ketamine/xylazine mixture to prevent movement-induced motoneuron activation, and then injected intraperitoneally with OCN. One hour later, mice were perfused with cold PBS, followed by cold 4% paraformaldehyde (PFA) in PBS. The lumbar spinal cord was post-fixed for 2 h in the same solution and transferred to 30% sucrose for cryoprotection at 4 °C. Spinal cords were embedded in Tissue-Tek O.C.T. (Sakura Finetek), cut into serial 20 μm sections (30 sections across 10 slides) on a cryostat, and stored at −20 °C.

For immunostaining, sections were blocked with 10% fetal bovine serum in TBS–0.15% Triton X-100 for 60 min at room temperature, and incubated overnight with rabbit anti-cFOS (1:500, Merck, ABE457), followed by Alexa-647 anti-rabbit and FluoroNissl Green to localize motoneurons. For each animal, 5–6 images were acquired using a Zeiss Spinning Disk fluorescence microscope. Motoneurons with nuclear cFOS staining were counted applying the same threshold across all groups, and averaged to obtain a single value per mouse, which was used as the statistical unit. All image acquisition and analysis were performed by a researcher blinded to the experimental group.

### GFP and SQSTM1/p62 immunostaining and puncta quantification

Mice were deeply anesthetized with a mixture Ketamine/Xylazine and transcardially perfused with cold PBS, followed by cold 4% Paraformaldehyde (PFA) in PBS. Post-fixation of spinal cords and cutting was performed as described above. For p62 immunostaining, sections were blocked with 10% fetal bovine serum for 60 min in TBS-Tx 0,15% at room temperature and then incubated with guinea pig anti-p62 (1:500, GP62C, Progen) overnight at 4°C. The sections were washed with TBS-Tx 0,3% before and after being incubated with an Cy3 Fluor-conjugated secondary antibody donkey anti-guinea pig IgG (1:500, Life Technologies) for 2h at room temperature in blocking buffer. Sections were incubated with Nissl Green (1:200, N21480, ThermoFisher), DAPI, and were mounted onto gelatin-subbed slides and cover slipped using Fluoromount. For p62 puncta analysis, images were obtained using a Zeiss Spinning Disk fluorescence microscope and the number of SQSTM1/p62 puncta was quantified on digital images with the spot detector in Icy software (http://icy.bioimageanalysis.org). The same threshold was applied uniformly across all images. For each animal, 5–6 images containing a total of 20–25 MNs were acquired. Puncta were counted within individual MN soma and normalized to soma area; values were averaged to obtain a single value per mouse, which was used as the statistical unit. All immunofluorescence image acquisition and analysis were performed by a researcher who was blinded to the experimental group.

For GFP staining, sections were blocked with 10% fetal bovine serum for 60 min in TBS-Tx 0,15% at room temperature and then incubated with rabbit anti-ChAT (1:500, PA5-29653, Invitrogen) overnight at 4°C. Then, the sections were washed with TBS-Tx 0,3% and incubated with Alexa555 against rabbit (1:500, Life Technologies) and with GFP-Booster (1:200, gb2AF488, ProteinTech) for 2h at room temperature, washed and mounted, and imaged using a Nanozoomer automated Slide Scanner.

### LC3b and HSP60 staining and quantification

NSC34 cells were differentiated and treated with OCN alone or in combination with BAF for 1 hours. For the immunostaining, coverslips were blocked with 10% fetal bovine serum for 60 min in TBS-Tx 0,15% at room temperature and then incubated with mouse anti-LC3b (1:500, M152-3, MBL) and rabbit anti-HSP60 (1:500, 12165S, Cell Signaling), overnight at 4°C. The sections were washed with TBS-Tx 0,3% before and after being incubated with an Alexa Fluor-conjugated secondary antibodies: donkey anti-rabbit IgG (1:500, Alexa Fluor 488, Life Technologies) and donkey anti-mouse IgG (1:500, Alexa Fluor 647, Life Technologies) for 2h at room temperature in blocking buffer. All sections were mounted onto gelatin-subbed slides and cover slipped using Fluoromount with DAPI, and 5-6 images per coverslip were acquired using a Zeiss Spinning Disk fluorescence microscope (Zen software, Zeiss). Colocalization analysis was performed in ImageJ/Fiji using the ComDet plugin (Spots Colocalization, https://imagej.net/plugins/spots-colocalization-comdet). Fold change was calculated to analyse whether OCN increases or decreases the presence of mitochondria in the autophagosome structures.

### Semi-quantitative RT-PCR

Spinal cord lumbar segments were immediately flash-frozen after dissection and total RNA was isolated with TRIzol Reagent using a homogenizer. Single-strand cDNA was synthesized from total RNA (1μg) by using SuperScript II Reverse Transcriptase. qRT–PCR was performed using iTAQ SYBR Green (BioRad). The following primers were used: *Gpr158* primers (5’- GCTCATCCTGTTGGAAACAAT-3’ and 5’- TGGCTCAAAGTACAGAATAACGAC-3’), *Pink1* primers (5’-TTCTTCCGCCAGTCGGTAG -3’ and 5’- CTGCTTCTCCTCGATCAGCC -3’), *Parkin2* primers (5’- GGAAATGGACCCTCCAATATGG -3’ and 5’- TCAAATCGCCCTTTCATAGCTTG -3’), *CREB1* primers (5’- AGCAGCTCATGCAACATCATC -3’ and 5’- AGTCCTTACAGGAAGACTGAACT -3’), *Bdnf* primers (5’- TCATACTTCGGTTGCATGAAGG -3’ and 5’- AGACCTCTCGAACCTGCCC -3’), *Beclin1* primers (5’- ATGGAGGGGTCTAAGGCGTC -3’ and 5’- TCCTCTCCTGAGTTAGCCTCT -3’), *Map1lc3b* primers (5’- TTATAGAGCGATACAAGGGGGAG -3’ and 5’- CGCCGTCTGATTATCTTGATGAG-3’), and *Gapdh* primers (5’- AGGTCGGTGTGAACGGAT -3’ and 5’- GGGGTCGTTGATGGCAACA -3’). All qPCR analyses were performed by an investigator blinded to treatment conditions.

### Pharmacological modulation of autophagy and mitophagy, and OCN treatment

*In vivo* pharmacological manipulation of targets or OCN treatment was performed by intrathecal injections of the following compounds: 100 ng of OCN, 10 µg of Urolithin A (SML1791, Sigma-Aldrich), 50 µg of Spautin-1 (SML0440, Sigma-Aldrich), 1 µg of Leupeptin (L2023, Sigma-Aldrich), 10 µg of Tat-Scramble or Tat-Beclin1 (Sigma-Aldrich) (REF), 20 µg of MTK-458 (7980, Tocris), in a total volume of 10 μL, administered 15 hours before the rotarod behavioral task. Mice were anaesthetized with a ketamine and xylazine cocktail as described before for the intrathecal injections. For the molecular studies, spinal cords were harvested 4 hours post-injection. All the drugs were diluted in NaCl. TETA was administered in the drinking water at 3% weight/volume (Castoldi et al., 2020; Pietrocola et al., 2020). Behavioural testing was performed 1 month after the start of TETA treatment.

### Recombinant OCN production

Mouse uncarboxylated OCN was purified from BL21 bacteria transformed with pGEX2TK-mOCN plasmid as previously described (Khrimian et al., 2017). In brief, GST-OCN fusion protein was bacterially produced in BL21 pLyS transformed with pGEX- 2TK-mOCN after induction with IPTG. Cells were collected in lysis buffer (PBS 1X 10 mM Tris, pH 7.2, 2 mM EDTA, 1% Triton, and 1X protease and phosphatase inhibitor cocktail; 78443; Thermo Fisher Scientific). Following four freeze-thaw cycles and sonication, lysates were cleared by centrifugation. Then, the supernatant was incubated with glutathione-Sepharose 4B (17075601; GE) during 4 hours at 4°C, washed it 6 times with washing buffer (PBS 1X and 1% Triton), and finally with PBS 1X. OCN was then cleaved out from the GST moiety by using thrombin (27-0846-01; GE). Four fractions were collected, and each of them was incubated with benzamidine Sepharose (17-5123-10; GE) for 30 min at room temperature to remove thrombin.

### Electrophysiology in spinal cord explants

Animals were anaesthetized with an overdose of xylazine (50 mg/kg) and ketamine (500 mg/kg), and the spinal cord was quickly removed and placed in a Na⁺-free solution containing (in mM): KCl 3, NaH₂PO₄ 1, glucose 25, sucrose 230, CaCl₂ 0.8, MgCl₂ 8, NaHCO₃ 26. It was then embedded in 2% agar (cooled to 35 °C) and further cooled after immersion to reach solidification. The agar block containing the spinal cord was cut with a vibratome into 400 µm-thick slices in a K-gluconate-based solution containing (in mM): K-gluconate 130, KCl 15, EGTA 0.05, HEPES 20, glucose 25. Slices were then transferred to an aCSF solution containing (in mM): NaCl 135, KCl 2.5, NaH₂PO₄ 1, glucose 25, CaCl₂ 2, MgCl₂ 1, NaHCO₃ 26, ascorbic acid 0.4, pyruvate 2, at 30 °C for 30 min and then at room temperature. Recordings started immediately. The intracellular solution contained (in mM): K-gluconate 144, KCl 6, MgCl₂ 4.6, HEPES 10, EGTA 1, CaCl₂ 0.1, Na-ATP 4, Na-GTP 0.4. All bicarbonate-containing solutions were bubbled with 95% O₂ / 5% CO₂.

Whole-cell current-clamp recordings were performed on ventral horn neurons, identified by their location and soma size. Neuronal excitability was assessed using slow triangular current ramps (5 s ascending, 5 s descending); the maximal ramp amplitude was set so that action-potential firing was consistent and well established at the start of the experiment. For each neuron, the current–frequency (I–F) relationship was fitted by linear regression before and after bath application of OCN (10–100 ng/ml), and the OCN-induced change in maximal firing frequency (ΔFmax) was derived along the regression line. The change in resting membrane potential (ΔV) was measured between conditions. Input resistance was computed from the sub-threshold membrane potential response at the start of the triangular current injection, plotted as a function of the injected current.

### Seahorse

Metabolic flux analysis was performed using an extracellular flux analyzer (Seahorse XFe96, Agilent). NSC34 cells were seeded in XFe96 cell culture microplates and allowed to adhere. Then, we applied the differentiation protocol for 4 days as described before. Before the assay, cells were incubated for 1 hour at 37 C in a non-CO2 incubator in unbuffered XF assay media with OCN treatment. For the measurement of the oxygen consumption rate (OCR), the media was supplemented with 10 mM glucose, 2 mM glutamine, and 1 mM sodium pyruvate. The mitochondrial stress test was conducted by sequentially injecting 1 uM oligomycin, 1 uM FCCP, and 0.5 uM rotenone/antimycin A to determine basal respiration, ATP-linked respiration, and maximal respiratory capacity. For the measurement of the extracellular acidification rate (ECAR), cells were incubated in unbuffered media containing 2 mM glutamine without glucose. The glycolysis stress test was performed by sequential injections of 10 mM glucose, 1 uM oligomycin, and 50 mM 2-deoxyglucose (2-DG) to evaluate basal glycolysis, glycolytic capacity, and glycolytic reserve. Following the assays, the OCR and ECAR values were normalized to the protein concentration per well.

### Adeno-associated viruses expressing shRNA, MITO-SRAI and MITO-DREADD

Adeno-associated viruses (AAV) serotype rh10 expressing shRNA were purchased from VectorBuilder (Chicago, US) or from the Viral Vector Production Unit (UAB, Spain). The promoters used here were H1 (general), SYN1 (specific for mature neurons), and mir218 (motoneuron-specific). The following shRNAs were used: Scr (AAVrh10-H1-Scr, AAVrh10-SYN1-Scr, AAVrh10-mir218-Scr), *Gpr158* (AAVrh10-H1-sh*Gpr158* and AAVrh10-mir218-sh*Gpr158*), *Beclin1* (AAVrh10-SYN1-shBeclin1), and *Pink1* (AAVrh10-SYN1-shPink1). AAVs overexpressing mito-SRAI (AAVrh10-SYN1-mito-SRAI) or mito-DREADD-Gs (Pagano Zottola et al., 2025)(AAVrh10-mir218-mito-DREADD) were purchased from VectorBuilder (US). All AAV titers were above 1.0×10^12^ GC/ml. AAVs were injected in a final volume of 10 μL (at least 1.0×10^10^ GC in total) 3 weeks before behavioural tests or spinal cord tissue collection.

### mito-SRAI analysis

Mice received an intrathecal injection of AAVrh10-SYN1-mito-SRAI and were allowed to recover. Three weeks later, vehicle or OCN was injected intrathecally. Four hours later, mice were transcardially perfused, and the spinal cord was harvested and sectioned on a cryostat (20 µm sections). Slices were washed and stained with FluoroNissl far-red to localize the MNs. For quantitative assessment of mito-SRAI signals, z-stack images were acquired using a Zeiss Spinning Disk fluorescence microscope (Zen software, Zeiss) and analyzed in Fiji/ImageJ (version 2.16.0/1.54p, National Institutes of Health, USA). The soma of the motoneurons was delineated based on FluoroNissl (far-red) staining. Once the region of interest (ROI) was defined, the fluorescence intensities of the two mito-SRAI channels (TOLLES and YPet) were measured within each ROI. The ratio between the blue (TOLLES) and green (YPet) channels was then calculated as the mitophagy index. Images were processed using uniform criteria to ensure comparability between groups. Images were acquired by one researcher, and the analysis was performed by another researcher blinded to the experimental group.

### Mitochondrial potentiation by Mito-DREADD

The mito-DREADD construct was kindly provided (Pagano Zottola et al., 2025), and used to generate an AAVrh10-mir218-mito-DREADD with VectorBuilder. Wild-type and *Gpr158*-/- mice received an intrathecal injection of AAVrh10-mir218-mito-DREADD and were allowed to recover. Three weeks later, vehicle or Clozapine N-oxide dihydrochloride (CNO) at 5 mg/kg (6329/10, Tocris) was injected intraperitoneally, and the rotarod test was performed 30 to 45 minutes later, as described for the novel object recognition test (Pagano Zottola et al., 2025). For molecular analysis, vehicle or CNO was administered, and spinal cords were harvested 45 minutes later.

### Behavioral tests

#### Rotarod

Locomotor function was assessed using an accelerating rotarod apparatus (Rota-rod/RS, Bioseb, France). Mice were placed on the rotating rod, with speed increasing progressively from an initial 4 rpm at a constant acceleration rate of 18 rpm/min, reaching a maximum of 40 rpm within 2 minutes; this maximum speed was then maintained for the remainder of the trial, up to a total trial duration of 5 minutes. The latency to fall from the rod was recorded for each trial. Testing was conducted over two consecutive days. On day 1, mice underwent a training phase consisting of four sessions per animal, each with a maximum duration of 5 minutes, to habituate them to the apparatus. On day 2, mice performed the test phase, consisting of five trials per animal under the same acceleration protocol, and the latency to fall was recorded for each trial.

#### Beam bar

Motor coordination and balance were further assessed using a graded bar beam test. The apparatus consisted of a 100 cm beam that tapered progressively in width from 3 cm to 1 cm, positioned horizontally between two cages. The end of the beam was made similar in appearance/bedding to the mice’s home cage to motivate the animals to cross the beam from the wide to the narrow end. Testing was conducted over two sessions: a training session, followed by a testing session, each consisting of three trials per animal. For each trial, the time taken to cross the beam, the total distance crossed and the distance when the 1st hindlimb slip happens were recorded as measures of motor coordination and balance.

### Open Field Test (OFT)

Mice were transported in their home cages over a short distance to the testing room and left undisturbed for at least 1 hour before testing. Each mouse was then individually placed in the center of a 43 × 43 cm open-field arena and allowed to explore freely during 30 minutes. Behaviour was monitored continuously using an infrared light beam activity monitor using actiMot2 Software (PhenoMaster Software, TSE System, Germany). General deambulatory activity was assessed by calculating the total distance travelled over the 30 minutes.

### Quantification and statistical analysis

All values are expressed as mean ± SEM. The alpha was set to 0.05 for all analyses. In each figure, the sample size and precise P values are included, and in the figure legend, the relevant statistical parameters and post hoc tests are reported in the corresponding legend. Group sizes were chosen based on pilot experiments and prior work employing comparable *in vivo* paradigms, in which 7 to 10 animals per group reliably produce robust and reproducible effects. The normality of each dataset was tested, and parametric tests (Student’s t-test or one-way ANOVA) or nonparametric alternatives (Mann–Whitney or Kruskal-Wallis) were selected accordingly. All analyses were performed in GraphPad Prism v9.

## Figure Legends Extented data

**Extended data figure 1: Gpr158 is expressed in the neuronal part of the motor system**

**a.-** Hierarchical clustering of GPCR expression (row-wise Z-score) across spinal cell types. Seven co-expression modules (Clusters 1–7) are indicated by the colour bar; Cluster 6, enriched in motoneurons (MNs) and containing Gpr158, is boxed. Red, expression above the gene mean across cell types; blue, below. **b.-** Global centrality change versus average logFC for GPCRs in each MN subtype, used to estimate the network impact of individual receptors. **c.-** Local network cluster (STRING) enrichment of Cluster 6 top genes, showing enrichment in family 3 / calcium-sensing GPCRs. FDR and gene count are indicated. **d.-** Relative mRNA levels of Gpr158 across the motor system relative to hippocampus (HpC), including: motor cortex, brainstem and spinal cord, and absent from skeletal muscle (soleus, gastrocnemius). **e.-** Representative Western blot of GPR158 in hippocampus, spinal cord, sciatic nerve, dorsal root ganglia and gastrocnemius. GAPDH was used as a loading control. **f.-** RNAscope in situ hybridization in the ventral horn of wild-type and *Gpr158*-/- spinal cord. *mGpr158* probe (green) combined with *mChat* for cholinergic neurons MNs (red) in WT, and is absent in *Gpr158*-/-spinal cord, confirming probe specificity. Scale bar = 50 µm.

**Extended data figure 2: Gpr158 downregulation in spinal cord MNs does not affect deambulatory activity**

**a-** Relative expression of *Gpr158* in the spinal cord of mice injected with AAVrh10-H1-Scramble or AAVrh10-H1-sh*Gpr158.* **b-** Representative fluorescent images of spinal cord cross-sections 3 weeks after injections with AAVrh10-mir218-Scramble expressing GFP. Spinal cord motoneurons were detected by ChAT staining (red). **c-** Relative expression of *Gpr158* in the spinal cord of mice injected with AAVrh10-H1-Scramble or AAVrh10-H1-sh*Gpr158.* **d-** Total distance in Beam bar test performed in wild-type and *Gpr158* -/- (male and female), and in male mice 3 weeks after intrathecal injections of AAVrh10-H1-Scramble, AAVrh10-mir218-Scramble, AAVrh10-mir218-sh*Gpr158*, AAVrh10-mir218-sh*Gpr158.* **e-** Open field test (OFT) performed in 3-month-old mice, 3 weeks after intrathecal injections with either AAVrh10-H1-Scramble, AAVrh10-mir218-Scramble, AAVrh10-H1-sh*Gpr158*, AAVrh10-mir218-sh*Gpr158*. For the OFT, distance and time spent in the centre of the arena (%), total ambulation (meter), and speed (cm/s) were measured.

Data are expressed as mean ± SEM. The P values were determined by Mann-Whitney test (**a, d, e**), Student’s t-test (**b, e**), or one-way ANOVA (Dunnett’s multiple comparison test) (**g**). Each dot represents an independent mouse (Spinal cord for molecular biology or behavioral test).

**Extended data figure 3: The osteocalcin–GPR158 axis controls motoneuronal gene expression, mirroring its signaling in the hippocampus**

**a-** Response of one motoneuron to a triangular injection of current in control conditions (left, blue trace) and after the application of OCN (10 ng/ml, right, red trace). The beginning of the response (< 1s, before firing starts) was used to compute the input resistance of the cell (by plotting the membrane potential as a function of the injected current). The bath application of OCN induced some changes to the input resistance (Δρ_ιν_ = -6.0 ± 20.5 MΩ, p=0.56, n=6, Wilcoxon test) and to the resting membrane potential (ΔV= 0.1 ± 5.4 mV, p=0.56, n=6 Wicoxon test). The variation of the resting membrane potential spans from -4.4 mV to +10.6 mV. The cell shown in (a) is one of the smallest reductions (ΔV= -2.2 mV). **b-** The difference between the maximum firing frequency during the ramps before and after OCN is plotted against the input resistance of the cell (before OCN). The size of the dot is proportional to the amplitude of ΔV (positive for the cells with an increased maximum firing frequency, negative for the cells that were inhibited). The purple dots represent two neurons that were exposed to a concentration of 100ng/ml of OCN. The open symbol corresponds to the cell shown in (a). **c-** Relative *ChAT* and *VAchT* expression in the lumbar spinal cord in wild-type (WT), *Ocn*-/- and *Gpr158*-/- mice, and 3 weeks after intrathecal injections with either AAVrh10-miR218-Scramble, AAVrh10-miR218-sh*Gpr158*. **d-** Relative *ChAT* expression in the lumbar spinal cord segment in wild-type after intrathecal injection of Vehicle or Osteocalcin (4h). **e-** Representative western blot and quantification analysis of GPR158 in the NSC34 cell line transfected with Scramble or sh*Gpr158*, and treated for 1 hour with vehicle or OCN. **f-** Relative *CREB* and *BDNF* expression in the lumbar spinal cord in wild-type (WT), *Ocn*-/- and *Gpr158*-/- mice. **g-** Total distance in beam bar test performed in wild-type and *Ocn* -/- (male and female).

Data are expressed as mean ± SEM. The P values were determined by Student’s t-test (**c**, **d**, **f**), one-way ANOVA (Dunnett’s multiple comparison test) (**e**) or Mann-Whitney test (**f**, **g**). Each dot represents an independent mouse (Spinal cord for molecular biology or behavioral test).

**Extended data figure 4: OCN and GPR158 regulates autophagy and mitochondrial proteins**

**a.-** Mean Z-scores of each protein cluster (C1–C8) across from wild-type (WT) intrathecally injected with Vehicle or OCN, and *Gpr158* -/- intrathecally injected with Veh, with pairwise statistical comparisons. **b.-** Proportional distribution of the regulated proteins across the three main categories (Category 1, 2 and 3). **c.-** Mean Z-scores of mitochondrial proteins across the 3 experimental groups. Several mitochondrial proteins are reduced in OCN-treated spinal cord, indicating that OCN induces mitophagy. **d.-** Ranking of all proteins by change in expression (delta Z) and network centrality. The core autophagy proteins Beclin1, Atg13 and Gabarapl1 emerge as the most central and most strongly regulated nodes**. e.-** Quantification of western blot of Beclin1 and LC3b-II in the lumbar spinal cord from wild-type WT, *Ocn-/-, Gpr158-/-,* upon intrathecal injections with AAV-miR218-Scramble (Scr) or AAV-miR218-shRNA*-shGpr158,* and upon intrathecal injection of Vehicle or OCN (related to **Fig. 4e**).

Data are expressed as mean ± SEM. The P values were determined by Student’s t-test (**e**) Each dot represents an independent mouse (Spinal cord for molecular biology).

**Extended data figure 5: Autophagy and mitophagy are required for osteocalcin/GPR158-dependent regulation of motor function**

**a-** Beam bar coordination test in 3-month wild-type (WT) and *Gpr158* -/- male mice systemically treated during 1 month with TETA. **b.-** Relative *Beclin1* expression in the lumbar spinal cord in wild-type mice 3 weeks after intrathecal injections with either AAVrh10-SYN1-Scramble or AAVrh10-SYN1-sh*Beclin1*. **c.-** Western blot quantification (related to **Fig. 5c**) of Beclin1, ChAT and LC3b in the lumbar spinal cord 3 weeks after intrathecal injections of AAVrh10-SYN1-Scramble and AAVrh10-SYN1-sh*Beclin1*. **d.-** Open field test (OFT) performed in 3-month-old mice, 3 weeks after intrathecal injections with either AAVrh10-SYN1-Scramble, AAVrh10-SYN1-sh*Beclin1*. **e-** Relative *Pink1* and *Parkin2* expression in the lumbar spinal cord in wild-type (WT), *Ocn*-/- and *Gpr158*-/- mice, and 3 weeks after intrathecal injections with either AAVrh10-miR218-Scramble, AAVrh10-miR218-sh*Gpr158*. **f.-** Relative *Pink1* expression in the lumbar spinal cord in wild-type 3 weeks after intrathecal injections with either AAVrh10-SYN1-Scramble or AAVrh10-SYN1-sh*Pink1*. **g.-** Open field test (OFT) performed in 3-month-old mice, 3 weeks after intrathecal injections with either AAVrh10-SYN1-Scramble, AAVrh10-SYN1-sh*Pink1*.

Data are expressed as mean ± SEM. The P values were determined by Student’s t-test (**b**, c, **d**, **e, f, g**), Mann-Whitney test (**c**, **d**, **e**), one-way ANOVA (Tukey’s multiple comparisons test) (**a**) or Kruskal-Wallis test (Dunn’s multiple comparison) (**a**). Each dot represents an independent mouse (Spinal cord for molecular biology or behavioral test).

**Extended data figure 6: Osteocalcin remodels the spinal cord metabolome:**

**a.-** Principal component analysis (PCA) of targeted metabolomics from lumbar spinal cord of mice 1 h after intrathecal injection of vehicle (Veh) or osteocalcin (OCN). **b.-** Heatmap of metabolite levels (row-wise Z-score) in Veh and OCN groups. **c.-** Western blot of pPka and PKA levels in *Gpr158-/-* lumbar spinal cord 3 weeks after AAVrh10-miR218-mitoDREADD injection and 45 mins after the injection of Veh or CNO. β-actin was used as a loading control.

Each dot represents an independent mouse (Spinal cord).

**Extended data figure 7: Aged mouse display motor defycits.**

**a.-** Rotarod and Beam bar coordination test in 3- and 16-month-old wild type mice. **b.-** Quantification of western blot against ChAT, Beclin-1, and LC3b-II in the lumbar spinal cord from 3- and 16-month-old mice (Relative to 7c). **c, d, e.-** Beam bar total distance test in the experimental groups from Figure 7e, g, and h.

Data are expressed as mean ± SEM. The P values were determined by Student’s t-test (**a**, **b**), Mann-Whitney test (**a**), or Kruskal-Wallis test (Dunn’s multiple comparison) (**c, d, e**). Each dot represents an independent mouse (Spinal cord for molecular biology or behavioral test).

## Acknowledgements and Funding

We thank the Imaging, Proteomics, Metabolomics, Histology, Behavioral, and Animal House Facility platforms of SFR Necker, and of the Animal House Facility of the BioMedTech Facilities, INSERM US36. This work was supported by HFSP (F.O.), ANR (CiliophAging) (F.O.), the ATIP-AVENIR Program (F.O.), AFM-Telethon Grant (F.O. and D.R.G), AFM-Telethon Trampoline Grant (D.R.G.), MSCA Fellowship (D.R.G), and École Doctorale (T.T and C.A.P)

## Author contributions

D.R.G. designed the experiments. D.R.G., T.T., A.K.T, S.M, B.L., C.A.P, S.D, I.N, S.M, performed the experiments. S.S. and D.M. performed the bioinformatic analyses. D.R.G. and T.T. analysed and interpreted the data. D.R.G and F.O. conceived the study. D.R.G. wrote the manuscript with input from all authors.

## Ethics statement

All behavioral experiments were performed in accordance with the European Communities for Experimental animal use (2010/63/EU) and local ethical committee review procedures and protocols.

