## Supplementary figures and images for "Bone regulates locomotion by sustaining motoneuronal mitochondrial function"

### Supp Figures

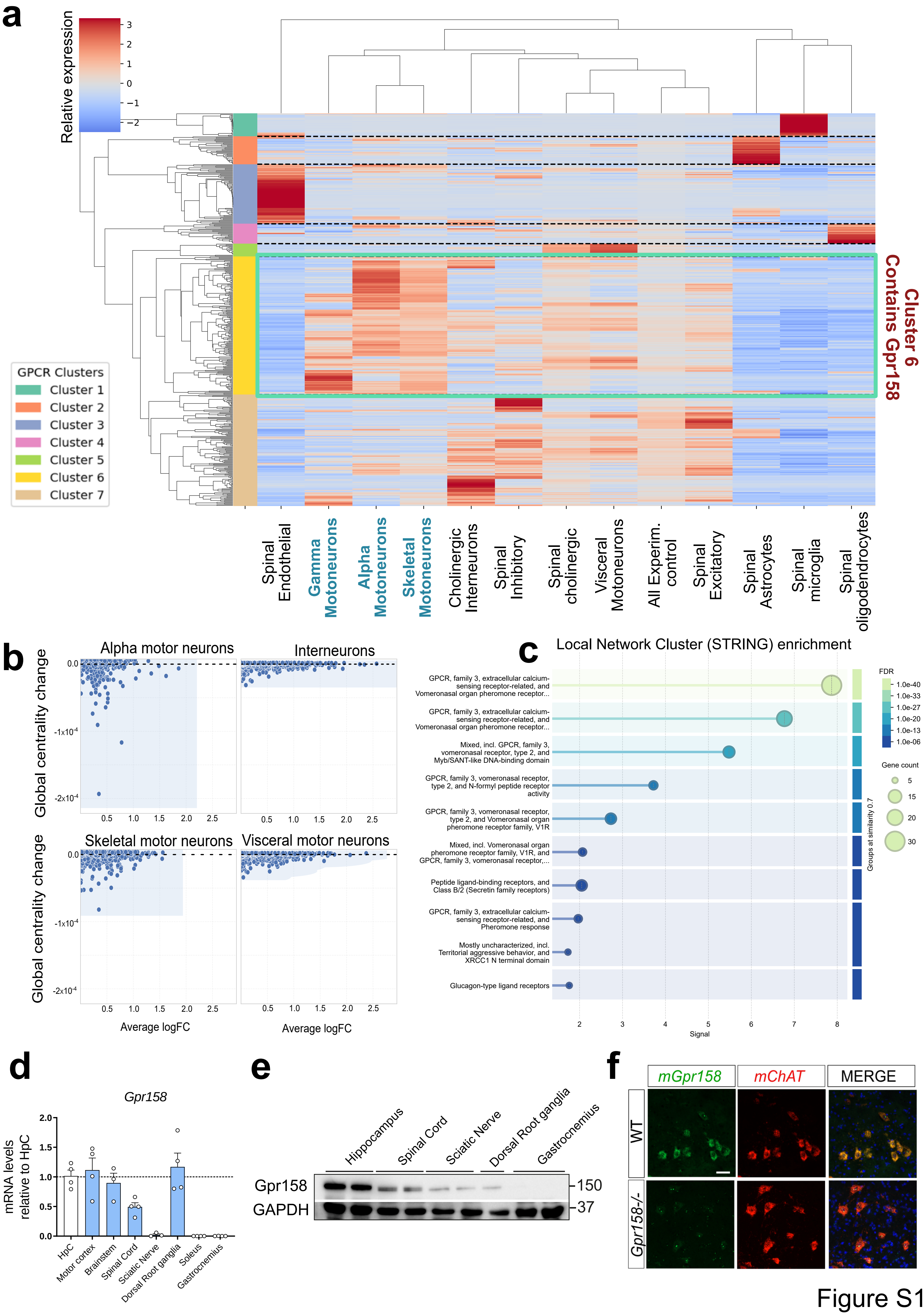

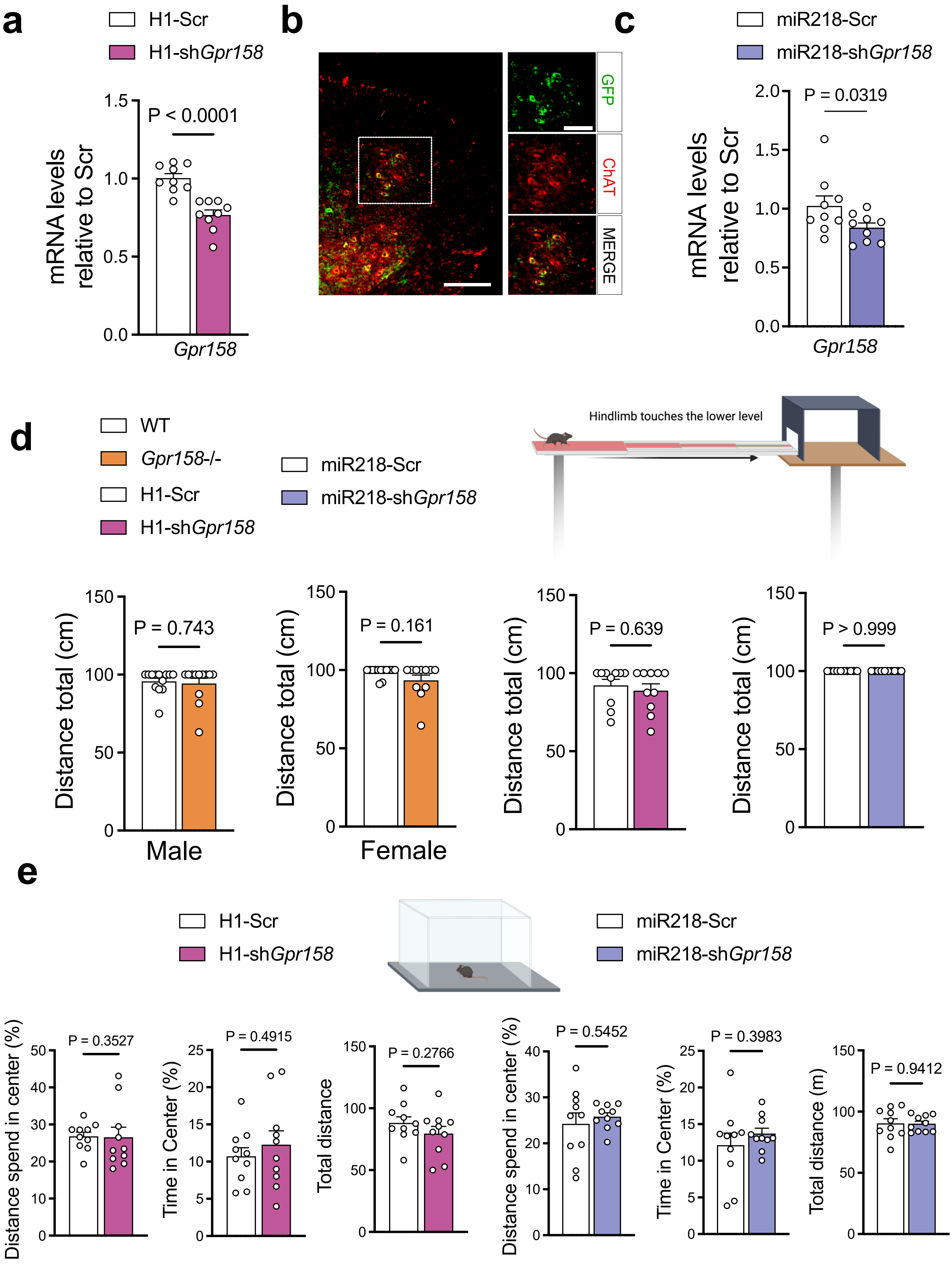

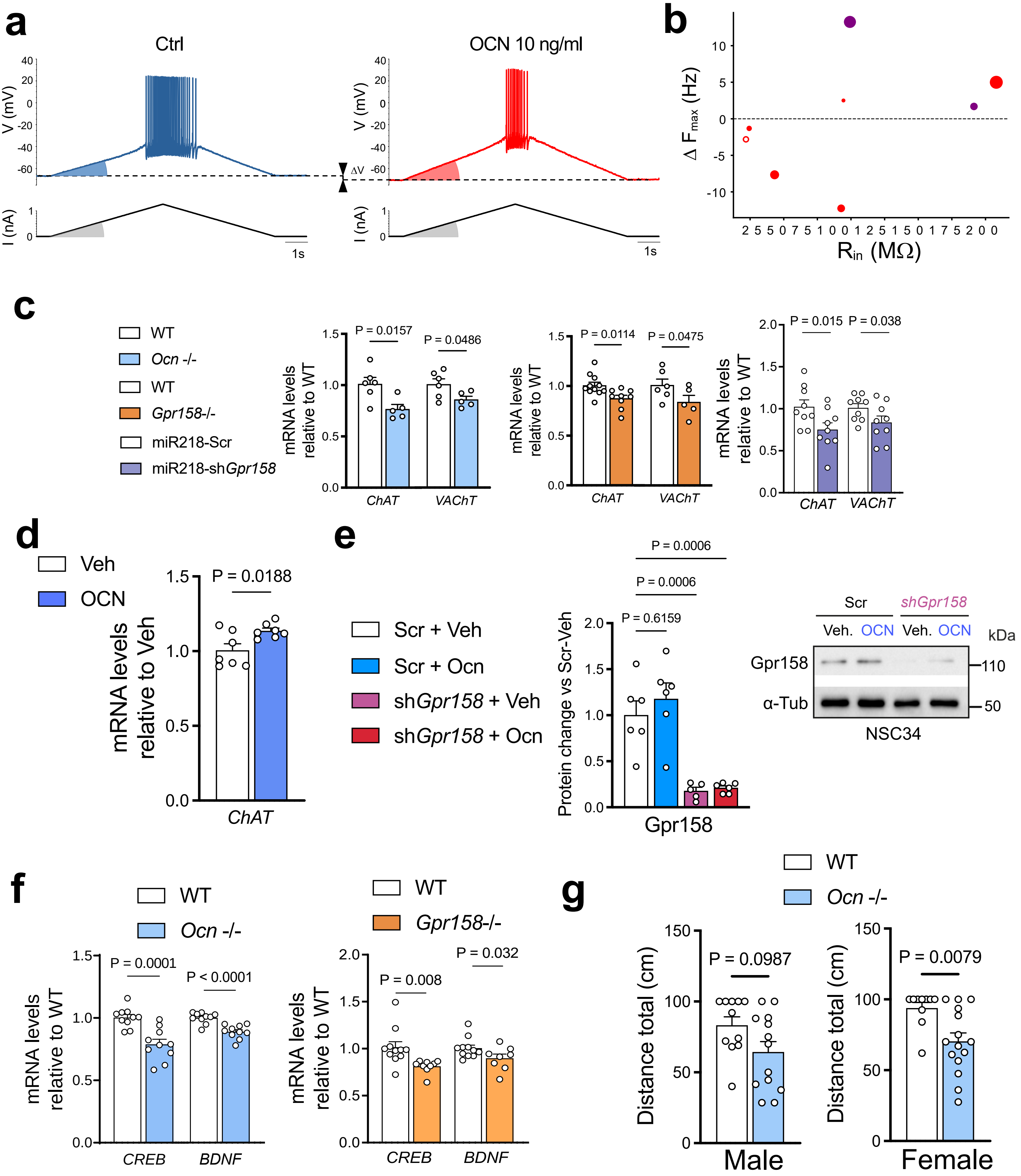

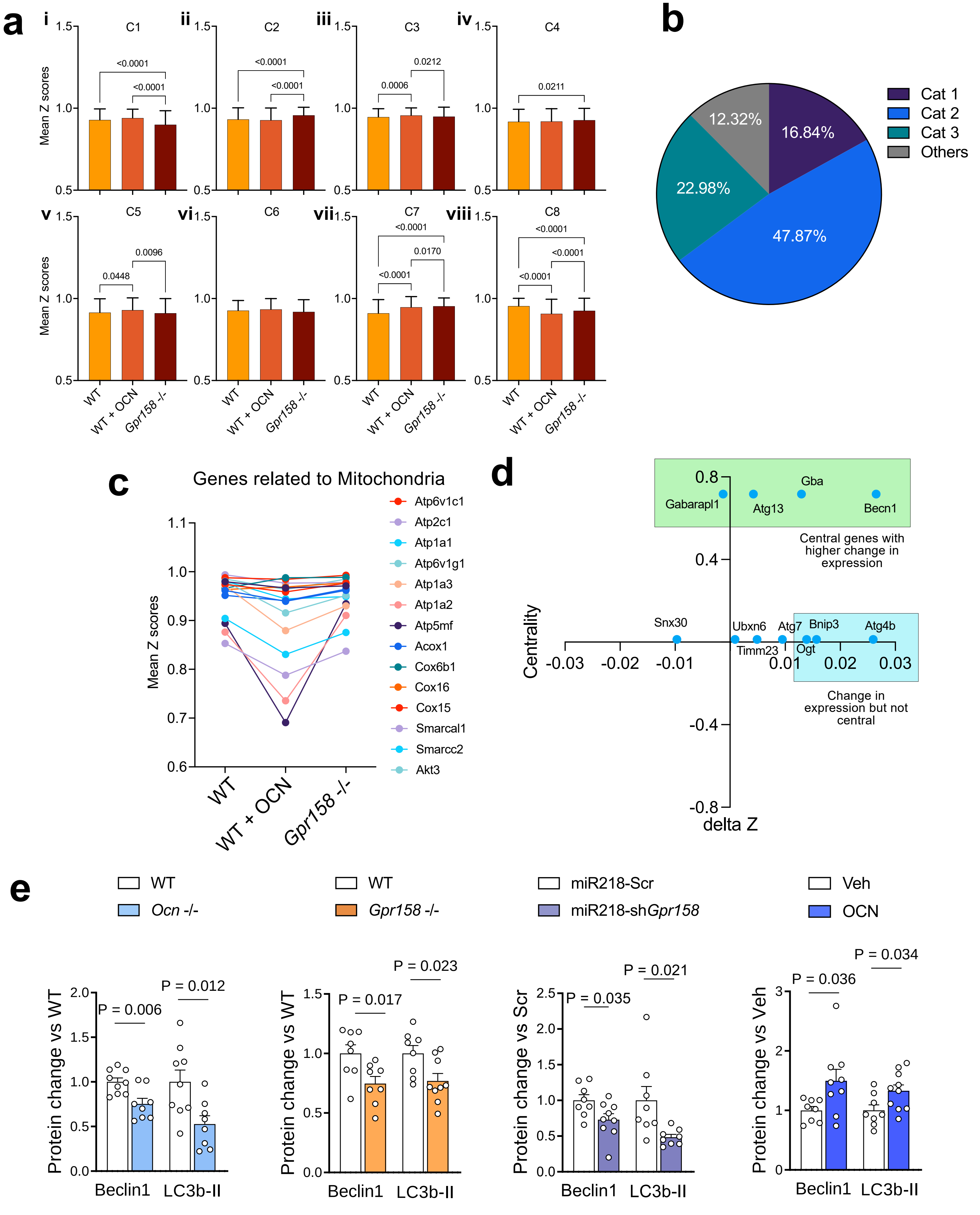

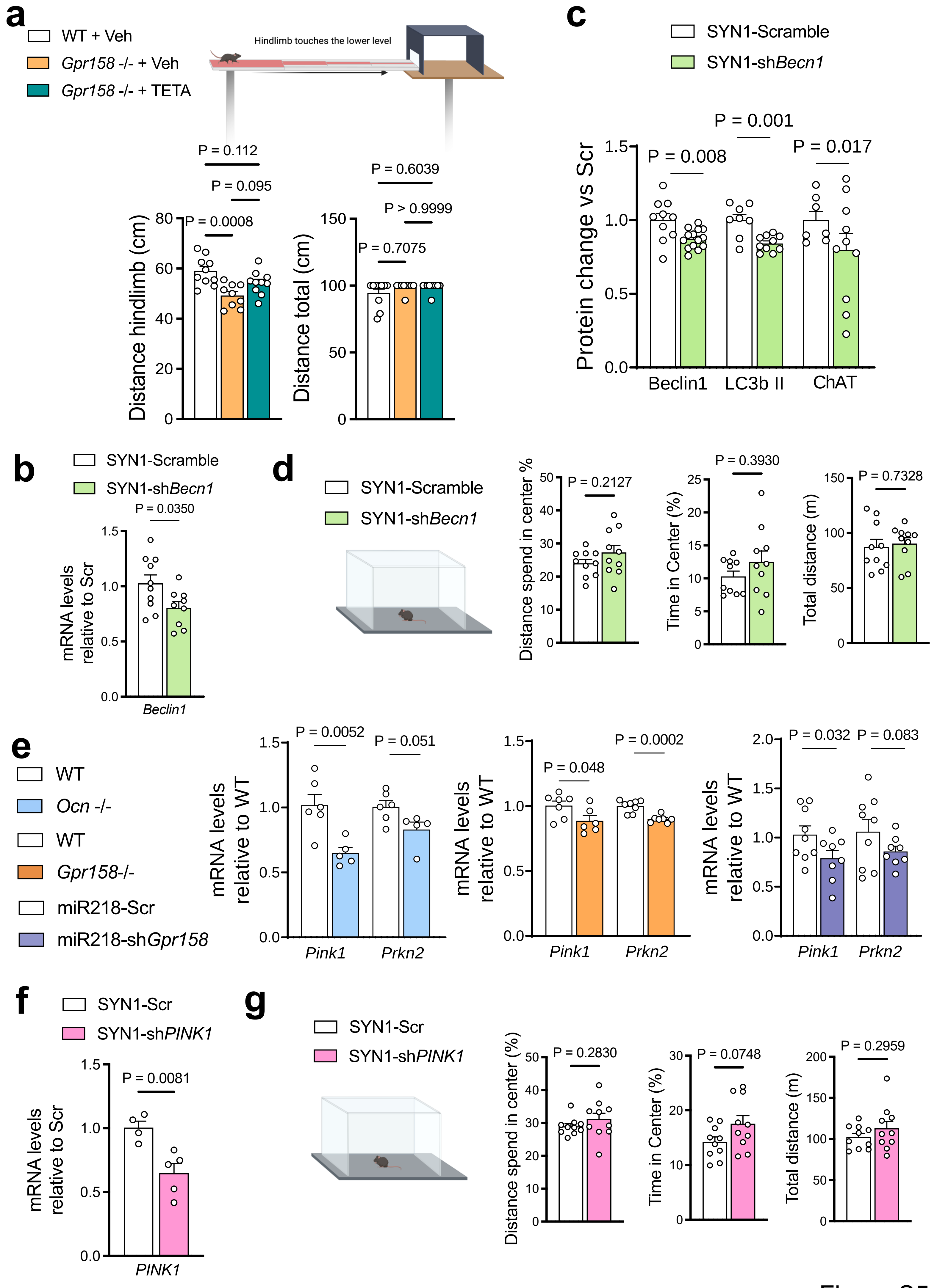

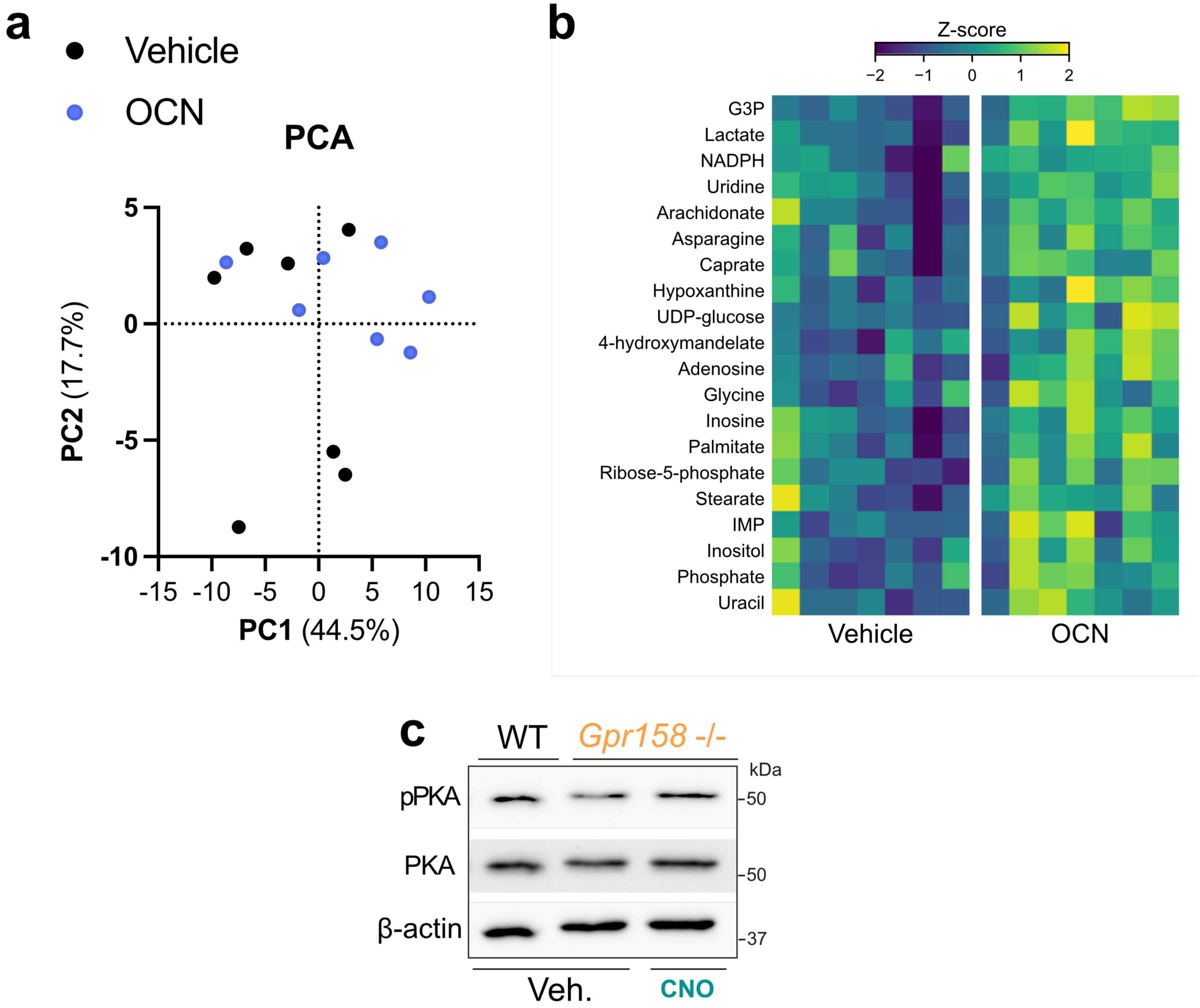

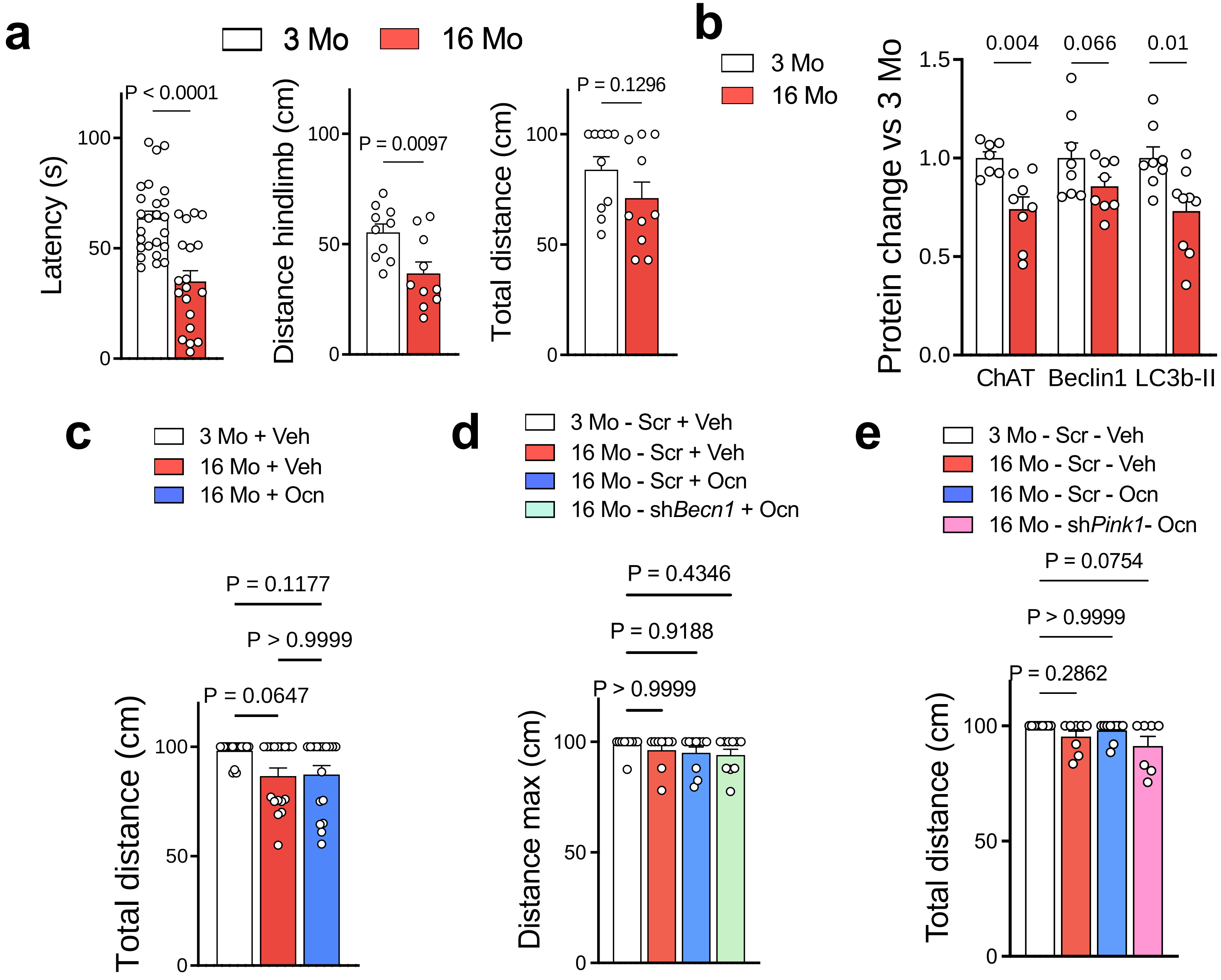
